# TLR2 Regulates Hair Follicle Cycle and Regeneration via BMP Signaling

**DOI:** 10.1101/2023.08.14.553236

**Authors:** Luyang Xiong, Irina Zhevlakova, Xiaoxia Z. West, Detao Gao, Rakhylia Murtazina, Anthony Horak, J. Mark Brown, Iuliia Molokotina, Eugene A. Podrez, Tatiana V. Byzova

**Affiliations:** Department of Neurosciences, Lerner Research Institute, Cleveland Clinic; Cleveland, OH 44195, USA; Department of Inflammation and Immunity, Lerner Research Institute, Cleveland Clinic; Cleveland, OH 44195, USA; Department of Cardiovascular & Metabolic Sciences, Lerner Research Institute, Cleveland Clinic; Cleveland, OH 44195, USA; Department of Biochemistry and Molecular Genetics, University of Illinois; Chicago, IL 60607, USA

## Abstract

The etiology of hair loss remains enigmatic, and current remedies remain inadequate. Transcriptome analysis of aging hair follicles uncovered changes in immune pathways, including Toll-like receptors (TLRs). Our findings demonstrate that the maintenance of hair follicle homeostasis and the regeneration capacity after damage depends on TLR2 in hair follicle stem cells (HFSCs). In healthy hair follicles, TLR2 is expressed in a cycle-dependent manner and governs HFSCs activation by countering inhibitory BMP signaling. Hair follicles in aging and obesity exhibit a decrease in both TLR2 and its endogenous ligand carboxyethylpyrrole (CEP), a metabolite of polyunsaturated fatty acids. Administration of CEP stimulates hair regeneration through a TLR2-dependent mechanism. These results establish a novel connection between TLR2-mediated innate immunity and HFSC activation, which is pivotal to hair follicle health and the prevention of hair loss and provide new avenues for therapeutic intervention.

**Summary:** Hair follicle stem cells TLR2 is required for hair homeostasis and regeneration. While TLR2 stimulation by endogenous ligand promotes hair growth, reduction in TLR2 and its ligand in aging and obesity may diminish hair growth.

## Introduction

Hair follicles (HFs), represent one of the best examples of mini-organs with the ability to regenerate throughout life, which, in turn, relies on the proliferation and differentiation of HF stem cells (HFSCs) within hair bulge (Fuchs and Blau, 2020, Sakamoto et al., 2021). The cyclic renewal of HFs is orchestrated by the interplay between inhibitory and stimulatory signals (Plikus et al., 2011). Despite the immune privileged status of HFs, they have a unique microbiome and immune system, including resident macrophages and other immune cells (Fuchs and Blau, 2020, Bertolini et al., 2020, Paus et al., 2003). Components of the HF immune system have been implicated in regulating the HF cycle and its regeneration (Rahmani et al., 2020, Di Domizio et al., 2020). Given their exposure to pathogens, HFs are equipped with innate immune receptors, particularly Toll-like receptors (TLRs), which detect and respond to pathogens by stimulating the secretion of defensins (Di Domizio et al., 2020, Selleri et al., 2007).

Toll-like receptors (TLRs) play a key role in recognizing and responding to either pathogen- or damage-associated molecular patterns, mediating the cytokine response. However, the role of TLRs extends beyond this function, as they have been shown to directly promote tissue regeneration and homeostasis in multiple tissues, particularly in stem and progenitor cells. TLRs regulate hematopoietic and intestinal stem cell renewal, proliferation, and apoptosis (Nagai et al., 2006, Tomchuck et al., 2008). The role of innate immune responses in the tissue-healing benefits of stem cell therapy has been clearly demonstrated (Vagnozzi et al., 2020). Moreover, TLR activation is a critical component of the reprogramming or transdifferentiation of adult cells into pluripotency (Lee et al., 2012), emphasizing the close coordination between innate immunity, cell transformation, and regeneration.

Multiple reports connect altered HFs immunity to hair loss, including a breakdown of immune privilege in alopecia areata (Rahmani et al., 2020). Likewise, androgen, which is tightly linked to TLR activation, was shown to influence the innate immunity of HFs in androgenic alopecia (Sawaya, 2012). The decline of innate immunity processes due to aging or conditions like obesity is widely recognized and these conditions are causatively associated with hair thinning and loss. (Andersen et al., 2016, Shaw et al., 2013, Palmer and Kirkland, 2016, Ghanemi et al., 2020). Alopecia patients often have higher body weight index and weight compared to healthy individuals (Bakry et al., 2014). Increased body weight index is linked to more significant hair loss severity in adults (Goette and Odom, 1976) and a higher prevalence of hair disorders in children and adolescents (Mirmirani and Carpenter, 2014). Mouse models support these findings, showing that activation of innate immunity through pathogen signals might lead to alopecia (Shin et al., 2018) and that high-fat diets inducing obesity cause hair thinning through HFSC depletion (Morinaga et al., 2021).

Our previous studies have shown that activating endothelial TLR2 by endogenous ligands such as CEP (a product of PUFA oxidation) promotes wound healing and tumor angiogenesis (West et al., 2010). Deletion of TLR2 from endothelial cells reduces tumor size by diminishing its vasculature (McCoy et al., 2021). In wounded skin, endothelial TLR2 is crucial for tissue regeneration through increased proangiogenic cytokine secretion (Xiong et al., 2022b). Although PUFAs have been shown to benefit hair growth by extending the anagen phase and promoting cell proliferation and hair shaft elongation (Munkhbayar et al., 2016), the role of innate immunity and in particular, TLR2 in the hair follicle cycle remains unknown.

Using animal models and human cell lines, we show a new function of TLR2 in the hair follicle cycle in homeostasis and hair follicle regeneration in injury. Furthermore, we demonstrate that an endogenously produced PUFA metabolite CEP serves as a TLR2 ligand in the hair bulge, promoting hair regeneration and growth through TLR2. In conditions associated with hair loss, i.e. aging and obesity, both TLR2 and its ligand are substantially depleted in HFs.

## Results

### TLR2 in HF declines due to aging and obesity

To assess whether and how aging affects hair follicle innate immunity, we analyzed available RNA sequencing data of mouse HFSCs (Doles et al., 2012). Pathway analysis revealed that innate and adaptive immunity, as well as TLR signaling, were among the top dysregulated pathways (Fig. 1A). Notch, JAK-STAT, TGF-β, and Wnt, and other pathways essential for HF regeneration were also altered by aging (Fig. 1A). Notably, the level of Tlr2 mRNA in HFSCs of old mice was two-fold lower compared to young mice (Keyes et al., 2013). In addition, TLR2 at the protein level was substantially lower in 13 months old mice compared to 2 months old mice (Fig. 1B, 1C).

**Figure 1.**
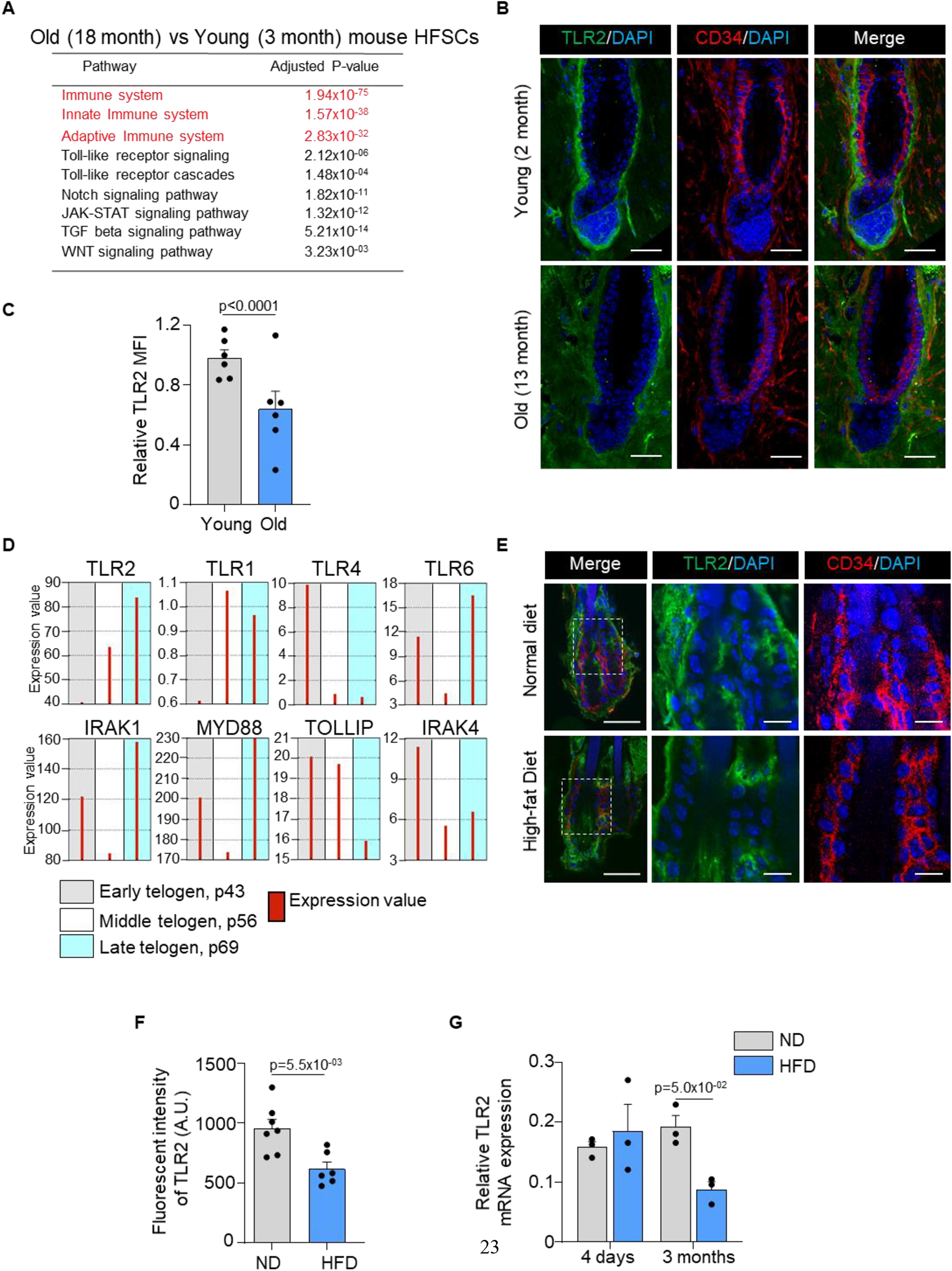
HFSCs downregulate TLR2 in response to stress like a high-fat diet and aging. A. Dysregulated pathways in old vs young mouse HFSCs. The top pathways are labeled red. B. Representative confocal images of telogen hair follicles from young and old mice immunostained for TLR2 and CD34 demonstrate decreased TLR2 intensity in HFSC (CD34-positive) of old mice. Scale bars are 50 μm. The middle and right panels show a magnified view of the boxed area. Scale bars are 20 μm. C. Quantification of TLR2 fluorescent intensity in images from B showing significantly lower TLR2 expression in HFSCs from the old mice. N =6 for each group. D. GEO2R analysis of published RNA data from sorted follicle populations in the 2nd telogen to anagen transition demonstrates the increased level of TLR2 mRNA accompanied by the activation of TLRs signaling downstream. E. Representative confocal images showing TLR2 expression in hair follicles from mice fed with a normal diet (ND) or high-fat diet (HFD). CD34 is an HFSC marker. Scale bars are 50 μm. Magnified images demonstrate decreased TLR2 intensity in HFSC (CD34-positive) of mice after HFD. Scale bars are 20 μm. F. Quantification of TLR2 fluorescent intensity in images from E showing significantly lower TLR2 expression in HFSCs from HFD-fed mice. N = 7 and 6 for ND and HFD groups respectively. AU, arbitrary unit. G. TLR2 mRNA expression in HFSCs from mice fed with ND or HFD for 4 days or 3 months. Data regenerated from published RNA sequencing dataset GSE131958. N = 3 for each group. All bar graphs are mean ± s.e.m. Non-parametric Mann-Whitney test (C, G) or unpaired two-tailed t-test (F) was used to determine statistical difference.

At the same time, the normal hair cycle is marked by an increase in TLR2 mRNA levels prior to HFSC activation during telogen, according to the analysis of existing RNA microarray data (Fig. 1D) (Greco et al., 2009). TLR2 mRNA levels are highest among other TLRs, with TLR1 and TLR4 mRNA showing the opposite pattern. TLR6 may act as a co-receptor for TLR2 as its expression pattern is similar. While the downstream TLR2 signaling molecules, IRAK1 and MYD88, are also upregulated, mirroring the TLR2, IRAK1 inhibitor TOLLIP is suppressed. Together, the results suggest the critical regulatory role of the entire TLR2/TLR6 pathway in the hair follicle cycle.

High-fat diet-induced obesity causes hair thinning and subsequent loss (Morinaga et al., 2021). In our model, a high-fat diet causes a nearly 2-fold decline in TLR2 protein level in HFSCs, compared to normal diet-fed mice (Fig. 1E, 1F). Further, RNA sequencing data reveal that 3 months of a high-fat diet is sufficient to reduce TLR2 levels in HFSCs by more than 2-fold compared to control mice (Fig. 1G) (Morinaga et al., 2021). Thus, our results and analyses of existing datasets demonstrate that conditions causatively associated with hair thinning and loss, such as aging and obesity, result in a dramatic depletion of TLR2 in HFSCs suggesting a possible regulatory role for TLR2 in hair follicles.

### TLR2 is upregulated during HFSC activation

The expression of TLR2 during a normal hair cycle was assessed using a previously characterized TLR2-GFP reporter mouse (Fig. 2), one of the best tools for the analysis of TLRs in vivo (Price et al., 2018). The correlation between the reporter and TLR2 protein expression was confirmed (Supplementary fig. 1A, 1B). The hair follicle cycle was verified by H&E staining (Supplementary fig. 1C). During telogen, a dormant stage for HFSCs, TLR2 was found in the bulge, secondary hair germ (sHG), dermal papilla (DP), and outer root sheath (ORS) (Fig. 2A). During regenerative anagen, TLR2 expression was detected in the DP and all sHG-derived progenitor cells (Fig. 2B, 2C) and in quiescent bulge stem cells (Fig. 2D, 2E). All cells derived from bulge stem cells (ORS) and sHG (hair shaft and inner root sheath, IRS) were positive for TLR2 (Fig. 2C). In catagen, TLR2 was abundant in the new bulge and sHG formed from ORS cells (Hsu et al., 2011) (Fig. 2F, 2G). While TLR2 was expressed in early sHG lineage (including IRS) (Fig. 2C), it was absent in mature (Fig. 2D, 2E) and regressing (Fig. 2F) IRS. This shows that TLR2 is abundant in stem cells but declines upon differentiation. The second telogen’s old and new bulges expressed TLR2 (Fig. 2H). TLR2 was present in sHG and DP during the second telogen (Fig. 2H) and increased in the late (competent) telogen compared to the early (refractory) telogen (Supplementary fig. 1D, 1E). The highest level of TLR2 occurred during active anagen compared to quiescent telogen and catagen (Fig. 2I). Quantitative PCR revealed that the Tlr2 mRNA level in HFSCs was 5- and 2.3-fold higher in anagen than in telogen and catagen, respectively (Fig.2J). Notably, analysis of existing RNA sequencing data using FACS-sorted cells (Lorz et al., 2010) confirmed that TLR2 expression was significantly higher in HFSCs than in epidermal or non-stem cells (Fig. 2K). Thus, TLR2 is enriched in HFSCs, and its expression increases during activation.

**Figure 2.**
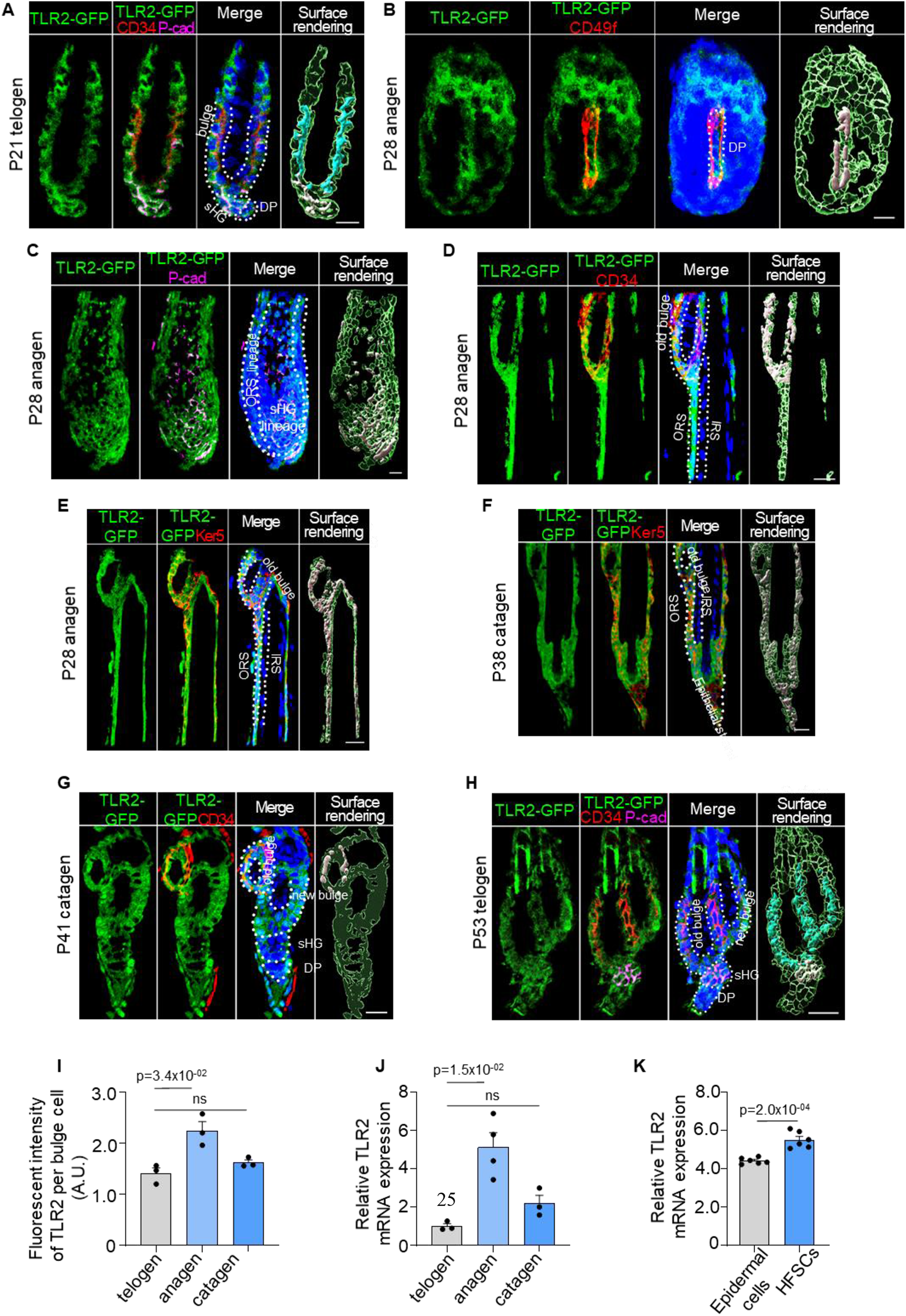
TLR2 is enriched in HFSCs and is upregulated during HFSC activation. TLR2-GFP reporter mouse skin sections were immunostained with anti-GFP to assess TLR2 expression in the hair follicles. A. Representative confocal images of P21 first telogen hair follicle immunostained for TLR2-GFP, CD34 (bulge stem cells), P-cad (secondary hair germ (sHG)), and DAPI (nuclei). The green color in the surface rendering panel represents TLR2 expression, and other surfaces show co-localization between TLR2 and specific markers. TLR2 is present in bulge, sHG, and dermal papilla (DP) cells. P represents postnatal days. Scale bar is 10 μm. B. TLR2-GFP in P28 anagen was co-immunostained with CD49f of basement membrane outlining the DP Scale bar is 10 μm. C. TLR2 is co-localized to the sHG lineage (P-cad^+^ layers), DP, and outer root sheath (ORS) lineage. Scale bar is 20 μm. D. TLR2-GFP in P28 anagen was co-immunostained with CD34 in old bulge (D) and Ker5 in ORS (E) revealing TLR2 localization to the old bulge, ORS, but not inner root sheath (IRS). Scale bars are 20 μm. F. Co-immunostaining of TLR2-GFP in P38 catagen hair follicle with Ker5 in ORS lineage cells showing co-localization of TLR2 with ORS and bulge. Scale bar is 20 μm. G. P41 late catagen hair follicle immunostained for TLR2 and CD34 showing co-localization of TLR2 to the old bulge, new bulge, sHG, and DP Scale bar is 20 μm. H. P53 second telogen hair follicle immunostained for TLR2, CD34, and P-cad reveals co-localization of TLR2 to the bulge, sHG, and DP Scale bar is 20 μm. I. Quantification of TLR2 fluorescent intensity in bulge cells at different phases showing TLR2 upregulation in anagen. N = 3 for each group. J. QPCR analysis of *Tlr2* mRNA expression in FACS-purified mouse HFSCs in anagen, telogen, and catagen. N = 3 or 4 per group. K. QPCR analysis of *Tlr2* mRNA expression in mouse epidermal cells and FACS-purified HFSCs showed significantly higher *Tlr2* expression in HFSCs compared with raw epidermal cells. N = 6 mice per group. All bar graphs are mean ± s.e.m. Two-tailed unpaired t-test (K) or Kruskal-Wallis test with Dunn’s post hoc test (I, J) was used to determine statistical difference.

### Deletion of TLR2 in HFSCs delays anagen onset in the normal hair cycle

To address the role of TLR2 in the hair cycle, we generated an HFSC-specific inducible Tlr2 knockout mouse line (TLR2^HFSC-KO^) and deleted Tlr2 during the first postnatal telogen (Xiong et al., 2022b). In TLR2^HFSC-KO^ mice, the telogen phase was substantially prolonged compared to control (TLR2^lox/lox^) mice, as summarized in the schematic (Fig. 3A). Melanogenesis and anagen onset are tightly coupled (Muller-Rover et al., 2001). Thus, the pink skin color at P21 marks the first postnatal telogen. The onset of anagen in control mice was indicated by the change in skin color to gray or black at P26 in control mice. TLR2^HFSC-KO^ mice at this age did not enter anagen, as evidenced by the delayed darkening of their skin (Fig. 3A-3C). This was further confirmed by skin section analysis at P21, P26, and P35 (Fig. 3D). At P26, control mice displayed pigmented anagen hair follicles with enlarged bulbs located deeply in the hypodermis, while nearly all follicles of TLR2^HFSC-KO^ mice were ∼5 fold shorter and remained in the dermis on top of adipose tissue, a characteristic of telogen. On day p35, we observe partial entrance into anagen in TLR2^HFSC-KO^ skin while the skin color of TLR2^HFSC-KO^ mice remains pink (Fig. 3D-3F). HFSCs were activated as early as at P24 in control mice based on positive Ki67 staining in sHG and bulge region (Fig. 3G), while most cells in TLR2^HFSC-KO^ sHG (Fig. 3H) and bulge (Fig. 3I) remained quiescent. At P25, control mice exhibited a large cluster of P-cad+ cells encapsulating DP within the transformed sHG (Fig.3K), whereas the sHG of TLR2^HFSC-KO^ mice remained small and inactive (Fig. 3J, 3K and Supplementary fig. 2). Despite the substantial delay in anagen onset, the morphology of hair follicles and expression of established HFSC markers, including Ker15, CD34, and Sox9, were normal in TLR2^HFSC-KO^ mice (Supplementary fig. 3). Thus, TLR2 in HFSCs is essential for HFSC activation and progression of the hair cycle.

**Figure 3.**
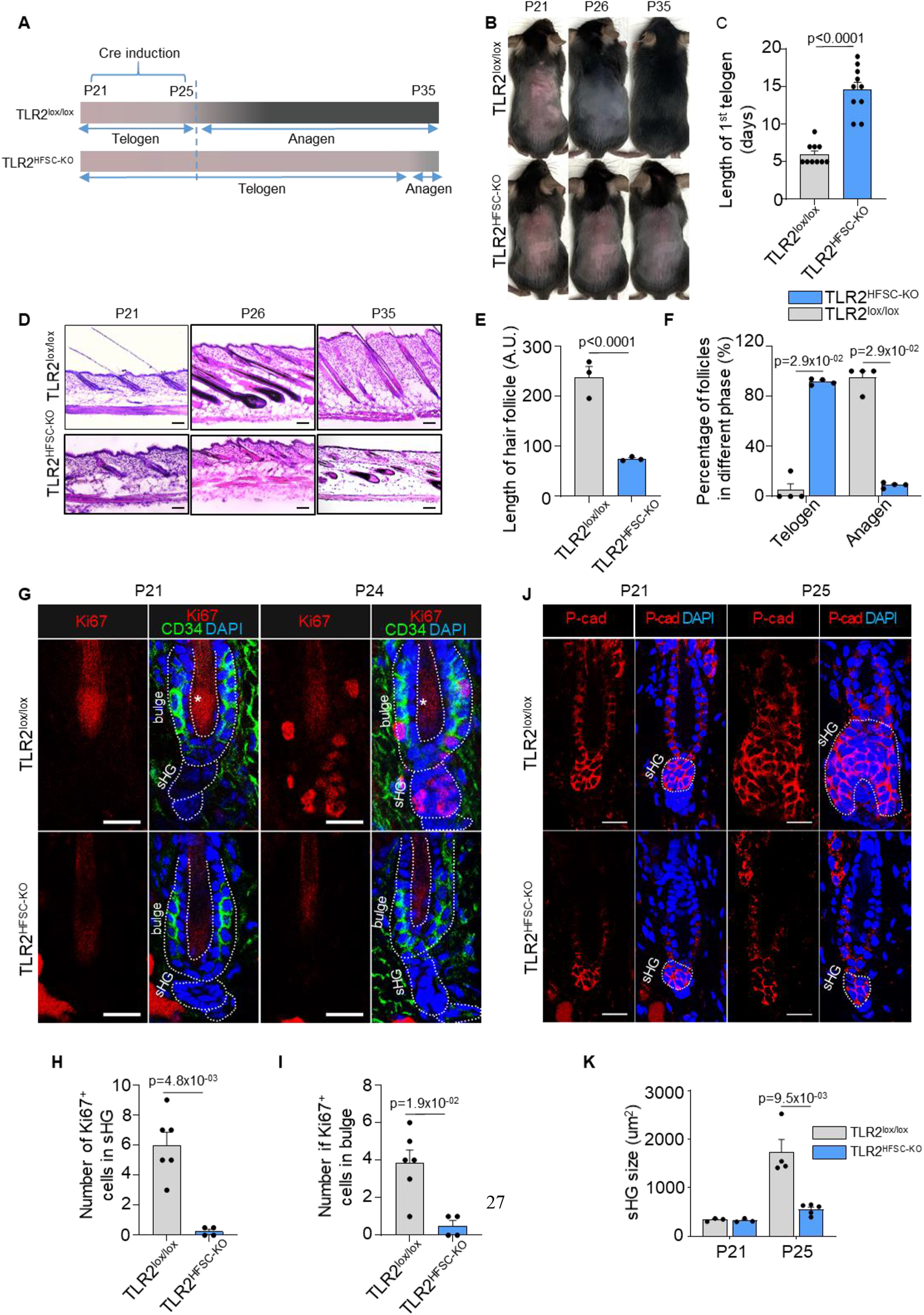
Deletion of TLR2 in hair follicle stem cells delays anagen onset. A. Schematic of RU486-mediated Cre induction and dorsal skin pigmentation change (gradient bars) in TLR2^lox/lox^ and TLR2^HFSC-KO^ mice. B. Representative images of shaved TLR2^lox/lox^ and TLR2^HFSC-KO^ mice showing different phases of the hair cycle. The TLR2^lox/lox^ mouse transitions from telogen (pink skin) to anagen (grey/black skin) at P26 and a full hair coat is developed by P35. The TLR2^HFSC-KO^ mouse exhibits a prolonged telogen (P21-P30-P35). Representative images of at least 10 mice in each group. C. Bar graph showing the length of first postnatal telogen starting from P21 measured by skin color change from B. N = 10 per group. D. Representative H&E staining of dorsal skin at indicated time points showing prolonged telogen in TLR2^HFSC-KO^ mice. Scale bars are 50 μm. E. The length of hair follicles at P26 from images in D. 50 hair follicles from 3 mice per group were used for quantification. F. Percentages of telogen or anagen hair follicles at P26 from D. N = 4 mice per group. G. Representative confocal images of P21 and P24 first telogen hair follicles from TLR2^lox/lox^ and TLR2^HFSC-KO^ mice immunostained for CD34, Ki67, and DAPI. Stars label the hair shaft. Scale bars are 20 μm. H, I. Quantification of images in G shows a diminished number of Ki67^+^ cells in sHG (H) and in CD34^+^ bulge (I) in TLR2^HFSC-KO^ mice compared to TLR2^lox/lox^ at P24. N = 6 and 4 mice for TLR2^lox/lox^ and TLR2^HFSC-KO^ group respectively. J. Representative confocal images of P21 and P25 dorsal skin sections from TLR2^lox/lox^ and TLR2^HFSC-KO^ mice immunostained for P-cad and DAPI showing changes in the size of sHG. Scale bars are 20 μm. K. Quantification of sHG size in panel K shows enlarged sHG in TLR2^lox/lox^ mice compared with TLR2^HFSC-KO^ mice. N=4 mice for p 25 TLR2^lox/lox^, and N=5 mice for TLR2^HFSC-KO^. Statistical significance was determined using a non-parametric Mann-Whitney test. All data are mean ± s.e.m.

### TLR2 regulates HFSC activation by interacting with BMP signaling pathway

The relationship between WNT and BMP signaling is central to the cyclic growth of hair follicles (Plikus et al., 2008). Anagen initiation is triggered by Wnt/β-catenin activation, while BMP signaling suppresses HFSC activation and its reduction is necessary for HFSC activation. Indeed, during the early (refractory) phase of the second telogen, HFSCs exhibit elevated BMP signaling as evidenced by high levels of BMP7 protein and pSMAD1/5/9, downstream targets of BMP signaling, compared to the late (competent) phase (Fig. 4A-4D). In contrast, BMP7 and its effectors, ID1 and ID2, are decreased during the late telogen based on our analysis of existing RNAseq datasets (Greco et al., 2009) (Supplementary Fig. 4A).

**Figure 4.**
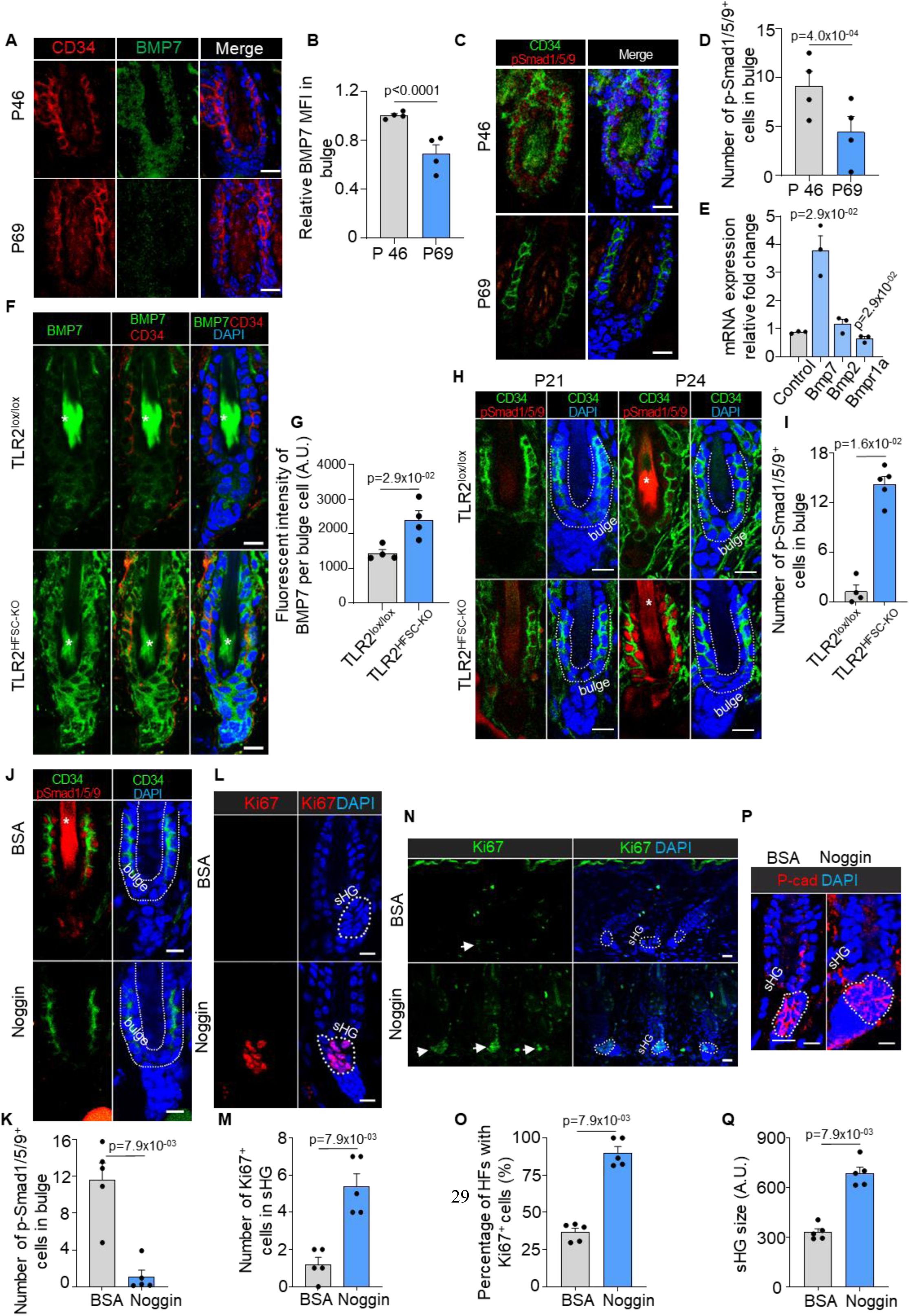
TLR2 interacts with BMP pathway to regulate the hair cycle. A. Representative confocal images of BMP7 staining in hair follicles of dorsal skin in early (p46) and late (p69) second telogen. Scale bars are 10 μm. B. Quantification of BMP7 fluorescent intensity from A showing diminished BMP7 expression during the second telogen from the early to late phases. N = 4 per group. C. Representative confocal images of pSMAD1/5/9 staining in hair follicles of dorsal skin in early (p46) and late (p69) second telogen. Scale bars are 10 μm. D. Quantification of pSMAD1/5/9^+^ positive cells in CD34^+^ bulge stem cells demonstrates a decrease of pSmad1/5/9 expression in late telogen. N = 4 per group. E. QPCR analysis reveals dysregulation of BMP singling molecules in HFSCs lacking TLR2. N=4 mice for Control and BMP2, N=3 mice for BMP7 and BMPr1A. F. Representative confocal images of BMP7 staining in hair follicles from TLR2^lox/lox^ or TLR2^HFSC-KO^ mice. Scale bars are 10 μm. Stars label hair shaft. G. Quantification of BMP7 fluorescent intensity from F showing higher BMP7 expression in TLR2^HFSC-KO^ mice. N = 4 per group. H. P21 and P24 dorsal skin sections from TLR2^lox/lox^ and TLR2^HFSC-KO^ mice immunostained for CD34, pSmad1/5/9, and DAPI. Scale bars are 10 μm. I. Quantification of pSmad1/5/9^+^ cells in CD34^+^ bulge stem cells in P24 dorsal skin from H. N = 4 and 5 for TLR2^lox/lox^ and TLR2^HFSC-KO^ respectively. J. Representative confocal images of dorsal skin sections from TLR2^HFSC-KO^ mice treated with BSA or noggin immunostained for CD34, pSmad1/5/9, and DAPI. Star labels the hair shaft. Scale bars are 10 μm. K. Quantification of pSmad1/5/9^+^ cells in CD34^+^ bulge stem cells from images in J. N = 5 per group. L. Immunostaining for Ki67 and DAPI in dorsal skin sections from TLR2^HFSC-KO^ mice treated with BSA or noggin. Scale bars are 10 μm. M. Quantification of images in L showing an increase in Ki67^+^ cells in sHG of noggin-treated compared to BSA-treated TLR2^HFSC-KO^ dorsal skin. N = 5 per group. N. Representative confocal images of Ki67 and DAPI immunostaining of dorsal skin sections from TLR2^HFSC-KO^ mice treated with BSA or noggin. Arrows point to hair follicles with Ki67^+^ cells in the sHG. Scale bars are 20 μm. O. Quantification of images in N showing percentages of hair follicles with Ki67^+^ cells in sHG. N = 5 per group. P. BSA- or noggin-treated TLR2^HFSC-KO^ mouse dorsal skin immunostained for P-cad and DAPI. The dashed line outlines the sHG. Scale bars are 10 μm. Q. Bar graph showing significantly larger sHG in noggin-treated TLR2^HFSC-KO^ mice. N = 5 per group. Mann-Whitney test was used to determine the statistical significance. All data are mean ± s.e.m.

To assess the possible connection between the TLR2 signaling and BMP pathway in human cells, we activated TLR2 and BMP signaling in human epidermal keratinocytes (NHEK) using a canonical TLR2 agonist (Pam3CSK4) and BMP4, respectively. As anticipated, BMP4 promoted the phosphorylation of its downstream target SMAD1/5/9. However, simultaneous co-activation of TLR2 diminished BMP4 signaling (Supplementary fig. 4B, 4C).

Likewise, stimulation of human HFSC with canonical TLR2 agonist Pam3CSK4 promoted cell proliferation by 1.5-fold compared to controls. Notably, this effect was diminished in the presence of a TLR2-blocking antibody (Supplementary fig. 4D, 4E). These results reveal that TLR2 activation on human HFSC augments their proliferation.

To gain a deeper understanding of TLR2’s role in HFSC activation, we profiled the transcriptome of HFSCs lacking TLR2 expression (Supplementary fig. 4F). The results showed that TLR2 deletion dysregulated 486 genes, many of which were involved in both the hair cycle and innate immunity. The most affected pathways included innate immunity response and TLR2 signaling, with its downstream target NF-kappaB (Supplementary fig. 4G). This profile somewhat resembles changes observed in aging models (Fig. 1A).

### BMP pathway is altered in TLR2 KO HFSCs

Since TLR2 suppresses BMP signaling and promotes HFSC proliferation, we assessed whether the delayed anagen in TLR2^HFSC-KO^ mice might be associated with the BMP pathway. QPCR analysis reveals that several key components of the BMP pathway were dysregulated in HFSCs lacking TLR2 (Fig.4E). Among those, the most notable changes were observed for BMP7, which was upregulated by ∼4-fold in TLR2-null HFSCs compared to controls (Fig. 4E). This was substantiated by co-staining of tissue sections for BMP7 and CD34, which demonstrated a ∼2-fold increase in BMP7 on HFSCs of TLR2^HFSC-KO^ mice as compared to the control (Fig. 4F, 4G). Activation of BMP signaling was assessed by pSMAD1/5/9 positive staining in HF. Quantification revealed a ∼15-fold higher level of pSMAD1/5/9 in TLR2-null mice (TLR2^KO^) as compared to controls (wild type) (Supplementary fig. 4H,4I). The significant increase in BMP signaling observed was attributed to the absence of TLR2 in hair follicle stem cells (HFSCs). This was evidenced by a comparable 14-fold increase in BMP signaling in follicles of TLR2^HFSC-KO^ mice compared to control during the first postnatal telogen phase, thereby ensuring the preservation of follicles in the dormant telogen stage (as shown in Fig. 4H and 4I). Simultaneously, the Wnt signaling and β-catenin stabilization within HFSCs, known to trigger their activation (Deschene et al., 2014), remained unchanged between control and TLR2HFSC-KO mice (as shown in Supplementary Fig. 4J).

### BMP antagonist rescues defects caused by the lack of HFSCs TLR2

To demonstrate that an altered BMP pathway is, indeed, responsible for the phenotype of TLR2 knockout in HFSCs, we utilized intradermal injection of noggin, a well-known inhibitor of BMP signaling (Botchkarev et al., 2001, Botchkarev et al., 1999), to block the upregulated BMP signaling in TLR2^HFSC-KO^ mice. As a result, noggin injection diminished activation of BMP signaling by >10-fold in TLR2^HFSC-KO^ mice as assessed by pSMAD1/5/9 staining of HF (Fig. 4J, 4K). Moreover, noggin promoted activation of TLR2^HFSC-KO^ HFs while the HFs in BSA-treated TLR2^HFSC-KO^ mice remained quiescent. Noggin treatment of TLR2^HFSC-KO^ mice dramatically upregulated cell proliferation within sHG as evidenced by Ki67^+^ cells (Fig. 4L, 4M), promoting ∼2.5 fold increase in activated follicles (Fig. 4N, 4O), thereby contributing to nearly 2-fold larger sHG (Fig. 4P, 4Q) as compared to BSA-treated TLR2^HFSC-KO^ mice. Thus, curbing suppressive BMP signaling in TLR2^HFSC-KO^ mice can reactivate their HFs, demonstrating a causative connection between TLR2 and BMP pathways in the hair cycle.

### HFSC TLR2 governs hair regeneration upon injury

High expression of TLR2 and its critical role in HFSC activation during the hair cycle prompted us to test the role of HFSCs TLR2 in an injury model where cells are more likely to be exposed to TLR2 ligands. First, we compared TLR2 levels in HFs in wounded and healthy skin using TLR2-GFP reporter mouse (Fig. 5A). In healthy skin, HFSCs upregulated TLR2 during their transition from middle to late telogen (day 5 to day 10) (Fig. 5B upper panels, and grey bars in fig. 5C), consistent with RNA sequencing results (Greco et al., 2009). This increase in TLR2 precedes HFSCs activation during the normal cycle. However, in wound HFSCs, TLR2 was upregulated immediately after an injury resulting in 1.5-fold higher expression compared to normal unwounded skin (Fig. 5B, 5C).

**Figure 5.**
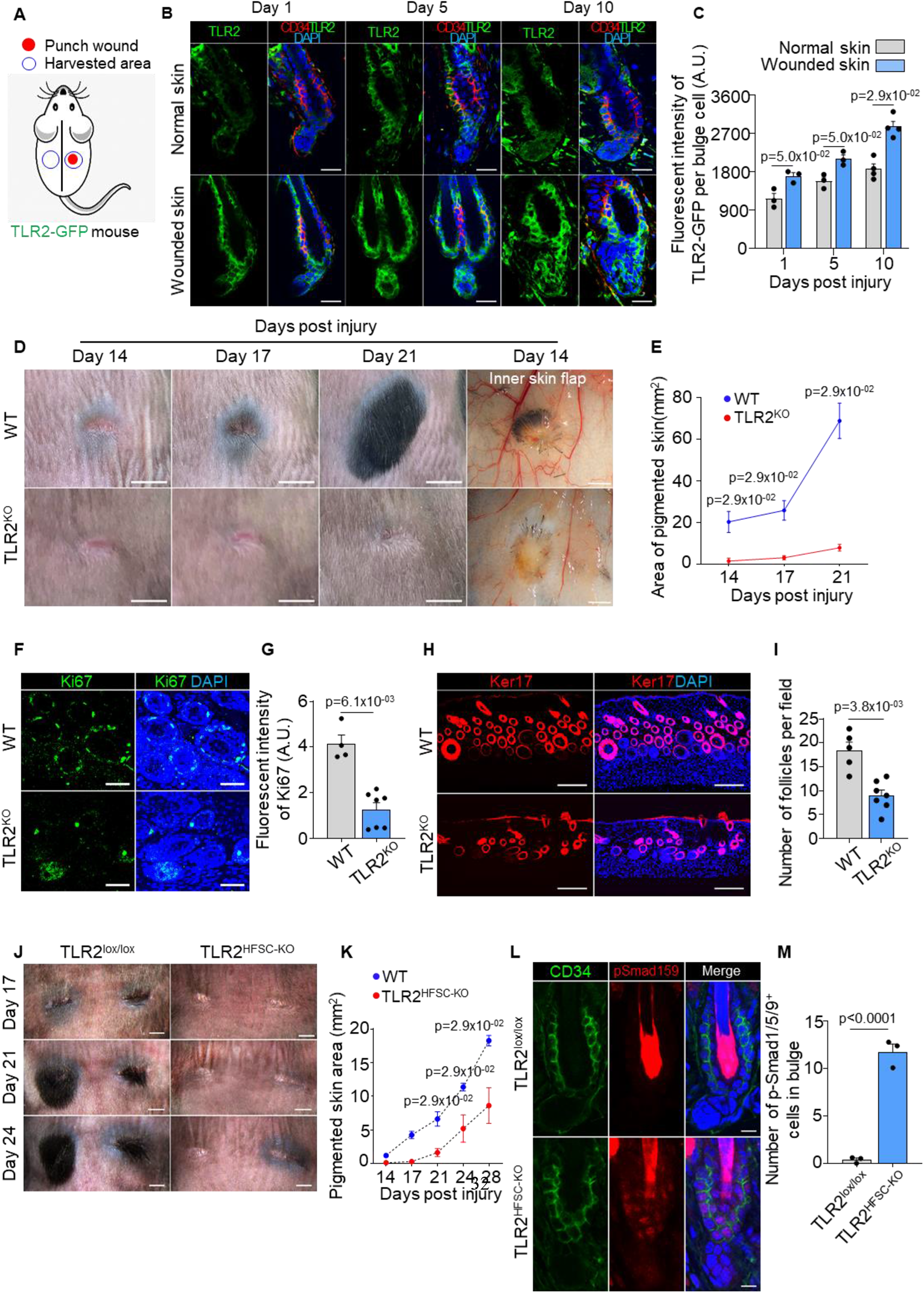
HFSC TLR2 is crucial for wound-induced hair follicle regeneration. A. Schematic of wound healing assay using TLR2-GFP reporter mouse. Full-thickness wounds on the dorsal skin of TLR2-GFP mice were created. Normal unwounded skin and the skin adjacent to the wound were harvested at different time points. B. Representative confocal images of normal and wounded skin from TLR2-GFP mice at different time points post-injury immunostained for TLR2-GFP and CD34. Scale bars are 20 μm. C. Quantification of TLR2 fluorescent intensity per bulge cell in hair follicles from B shows increased TLR2 level in hair follicles from wounded skin as compared to normal skin. N=3 for Day 1 and Day 5, and N=4 for Day 10 per group. D. Representative photographs showing hair regeneration on the dorsal skin and inner skin flaps at indicated time post-injury in WT and TLR2 global knockout (TLR2^KO^) mice. Diminished hair growth around the wound is apparent in TLR2^KO^ skin from day 14 through 21 post-injury compared to WT skin. The inner skin flaps from TLR2^KO^ at day 14 post-injury show an absence of pigmented hair bulbs and skin pigmentation. Scale bars are 5mm for dorsal skin and 1mm for inner skin flaps. E. Quantification of pigmented dorsal skin area around the wound from images in D shows diminished pigmentation in TLR2^KO^ skin compared with WT skin at all time points post-injury. N=4 per group. F. Representative confocal images of skin adjacent to wound immunostained for Ki67 and DAPI. Scale bars are 50 μm. G. Bar graph showing diminished Ki67 fluorescent intensity in the skin adjacent to wound in TLR2^KO^ mouse compared to WT mouse from images in F. N = 4 and 7 for WT and TLR2^KO^ respectively. H. Representative confocal images of skin adjacent to wound immunostained for Ker17 and DAPI. Scale bars are 100 μm. I. Quantification of hair follicle numbers from images in H reveals a significant decrease in regenerated hair follicles in TLR2^KO^ skin compared with WT skin. N = 5 and 7 for WT and TLR2^KO^ respectively. J. Representative photographs showing a lack of hair regeneration and skin pigmentation around the wound on the dorsal skin of TLR2^HFSC-KO^ mice compared with TLR2^lox/lox^ mice on day 17, day 21, and day 24 post-injury. Scale bars are 2mm. K. Quantification of pigmented skin area around the wound during 14-28 days post-injury showing significantly smaller pigmented skin area in TLR2^HFSC-KO^ mice compared with TLR2^lox/lox^ mice. N = 4 per group. L. Representative confocal images of wounded skin from TLR2^lox/lox^ and TLR2^HFSC-KO^ mice stained for CD34 and pSmad1/5/9. Scale bars are 10 μm. M. Quantification of images from L showing more pSmad1/5/9^+^ cells in TLR2^HFSC-KO^ wounded skin. N = 3 per group. Mann-Whitney test was used to determine the statistical significance. All data are mean ± s.e.m.

Hair regeneration after injury represents a substantial part of the healing process (Wang et al., 2017, Chen et al., 2015, Abbasi and Biernaskie, 2019). We next assessed the role of TLR2 using age- and gender-matched WT and TLR2^KO^ mice. The lack of TLR2 visibly impaired hair regeneration after wound healing (Fig.5D). On day 14 post-injury, HFs in WT mice entered precocious anagen judged by a spot of pigmented skin, which, on day 21 developed into a black hair patch (Fig. 5D). In contrast, the follicles of TLR2^KO^ mice remained quiescent lacking regenerated HFs around wounds even after 21 days post-injury (Fig. 5D, 5E). At this point, pigmented skin area in WT was ∼9-fold larger than in TLR2^KO^ mice. Skin flaps showed substantial pigmentation and growing hair bulbs around wounds in WT, indicative of active anagen. In contrast, the TLR2^KO^ skin flap was devoid of pigmentation, consistent with telogen (inner skin flap in fig. 5D). Ki67 staining confirmed an increase in HFs activation in WT but not in TLR2^KO^ skin (Fig. 5F, 5G). The resulting density of regenerated HFs based on Ker17 staining in WT was 2-fold higher than in TLR2^KO^ mice (Fig. 5H, 5I). Most importantly, this effect was dependent on TLR2, specifically on HFSCs, since TLR2^HFSC-KO^ mice exhibited a similar phenotype with a dramatic reduction in pigmentation and hair growth compared to control mice (Fig. 5J, 5K). The upregulation of pSmad1/5/9 in TLR2^HFSC-KO^ wounds compared to controls demonstrates that similar to the HF cycle scenario, increased BMP signaling might contribute to diminished HF regeneration (Fig. 5L, 5M). Thus, the TLR2-BMP axis in HFSCs governs HF regeneration after injury.

### Endogenous ligand promotes hair regeneration via TLR2 on HFSCs

One of the most important endogenous ligands for TLR2 is CEP, which is a naturally occurring product of PUFA oxidation shown to be accumulated during inflammation and wound healing (West et al., 2010, Xiong et al., 2022a). Healthy tissues are typically devoid of this product, which is mainly associated with inflammation and pathologies (Yakubenko et al., 2018). However, in contrast to other tissues, healthy HFs exhibited high levels of CEP accumulation (Fig. 6A). During anagen, CEP is present within the proximal part of the follicle, while in telogen the entire follicle is encased by this PUFA metabolite (Fig. 6B-6E). Generation of CEP from PUFA is directly aided by myeloperoxidase (MPO) (Yakubenko et al., 2018, Xiong et al., 2022a). MPO is present in abundance in sebaceous glands, possibly as a part of immune defense (Supplementary fig. 5A). Even more surprising, in contrast to other organs and tissues, CEP in HFs is substantially depleted with age (Fig. 6F, 6G), and this decline coincides with the reduction in the regenerative potential of HFs. This is likely due to decreased level of MPO during aging (Supplementary fig. 5B, 5C).

**Figure 6.**
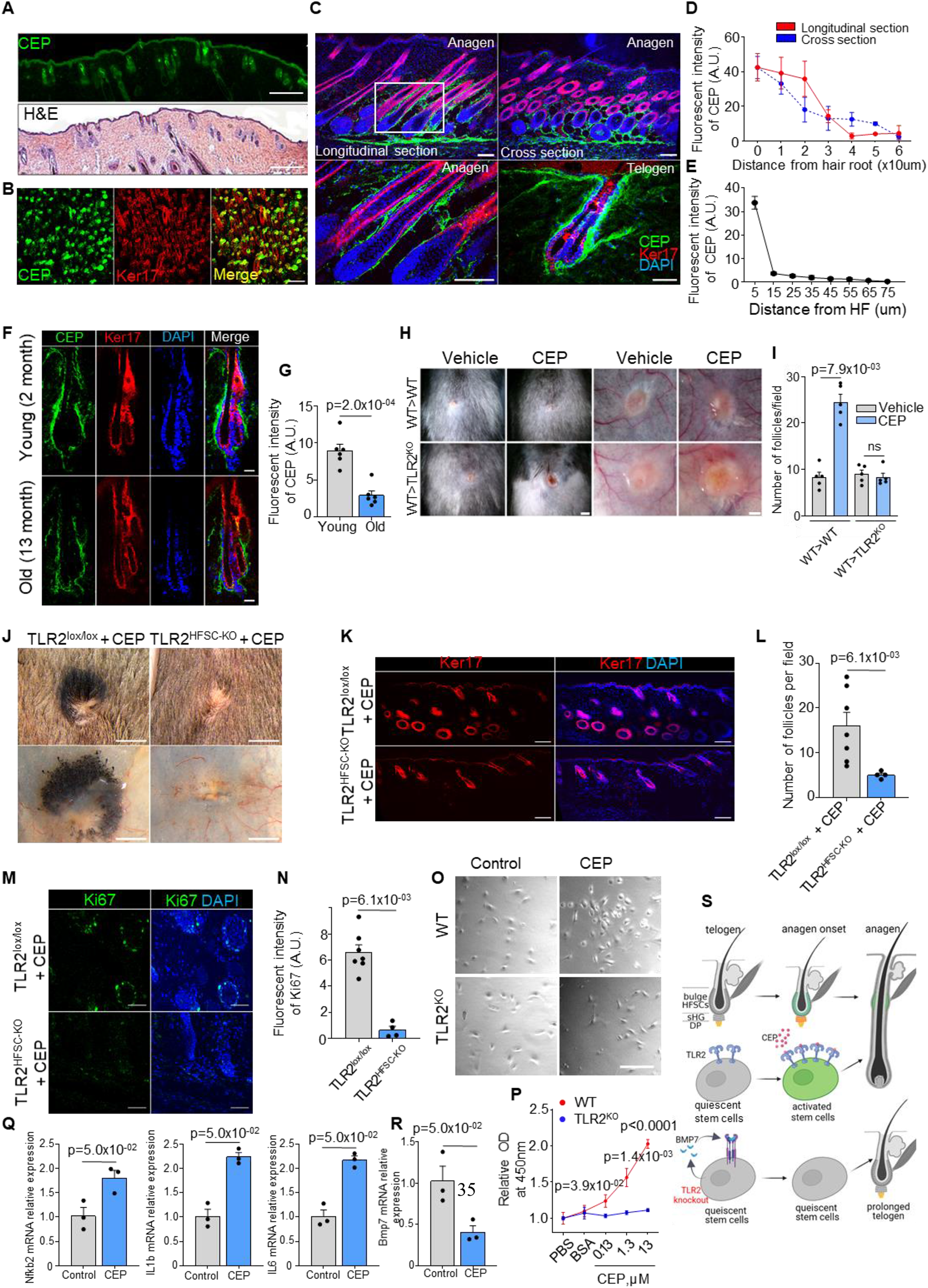
Oxidation-dependent TLR2 ligand CEP is present in hair follicles and promotes hair regeneration via HFSC TLR2. A. Representative images of H&E and CEP immunostaining of consecutive skin sections from WT anagen mouse. Scale bars are 1 mm. B. Representative confocal images of P5 WT whole-mount skin immunostained for CEP and Ker17. The merged image shows the co-localization of CEP to anagen hair follicles (Ker17^+^). Scale bar is 200 μm. C. Longitudinal and cross-sections of anagen and telogen hair follicles from WT mice immunostained for CEP and Ker17. The lower left panel shows a magnified view of the boxed area. Scale bars are 100 μm for anagen, 50 μm for telogen. D. Quantification of CEP fluorescent intensity at a different distance from the root of anagen hair follicles in longitudinal and cross-sections immunostaining images in images from C. A gradual decrease in CEP levels is observed from the proximal to the distal part of anagen hair follicles. N= 50 follicles from 3 per group. E. Line chart showing a sharp decrease of CEP fluorescent intensity with the distance from HF in telogen (from the lower right panel in C). N= 10 follicles from 3 per group. F. Representative confocal images of telogen hair follicles from young and old mice immunostained for CEP and Ker17. Scale bars are 20 μm. G. Quantification of CEP fluorescent intensity from images in F. N = 6 mice per group. H. Representative photographs of dorsal skin (two left panels) and inner skin flaps (two right panels) from WT and TLR2^KO^ mice after irradiation and bone-marrow transplantation of WT bone marrow demonstrate an increased number of pigmented hair bulbs and skin pigmentation around wounds in CEP-treated wounds compared to control in WT mice with no differences in TLR2^KO^ transplanted with WT bone marrow. Scale bars are 1 mm for the dorsal skin and 500 μm for the inner skin flap. I. Quantitative results from H show an increased density of hair follicles upon CEP application around wounds of WT> WT transplanted mice with no changes in WT> TLR2^KO^ mice. N = 5 for each group. J. Representative photographs of dorsal skin (upper panels) and inner skin flaps (lower panels) from TLR2^lox/lox^ and TLR2^HFSC-KO^ mice treated with CEP show a lack of pigmentation around TLR2^HFSC-KO^ wounds compared with TLR2^lox/lox^ wounds treated with CEP. The inner skin flap of TLR2^HFSC-KO^ mice demonstrates an absence of pigmented hair bulbs after the CEP treatment Scale bars are 3 mm. K. Representative confocal images of skin adjacent to wound immunostained for Ker17. Scale bars are 100 μm. L. Quantification of hair follicle numbers in images from K reveals a significant decrease in regenerated hair follicles in TLR2^HFSC-KO^ skin compared with TLR2^lox/lox^ skin. N=7 for TLR2^lox/lox^ N = 4 for TLR2^HFSC-KO^. M. Representative confocal images of skin adjacent to wound immunostained for Ki67. Scale bars are 50 μm. N. Bar graph showing Ki67 fluorescent intensity in the skin adjacent to wound from images in M. N=7 for TLR2^lox/lox^ N = 4 for TLR2^HFSC-KO^. O. Representative microphotographs of primary keratinocytes isolated from WT or TLR2^KO^ mouse skin co-cultured with CEP or control (PBS or BSA). Representative images from at least three independent assays are shown. Scale bar 50 µm. P. Cell proliferation of primary keratinocytes in O indicates increased proliferation by CEP in WT but not in TLR2^KO^ keratinocytes. N=3 independent experiments. Q. QPCR analyses of *Nfkb2*, *Il1b*, and *Il6* mRNA levels in FACS-purified mouse HFSCs treated with BSA control or CEP. N = 3 per group. R. QPCR analyses of *Bmp7* mRNA levels in FACS-purified mouse HFSCs treated with BSA control or CEP. N = 3 per group. S. Summary of the main findings of this study. Unpaired t-test (G, P) or Mann-Whitney test (I, L, N, Q, R) was used to determine the statistical significance. All data are mean ± s.e.m.

The connection between CEP levels and hair thinning and loss in aging prompted us to test whether exogenous CEP can activate TLR2 in HFSCs and stimulate their proliferation. Our in-vitro experiments revealed that CEP increases the proliferation of human HFSC in a TLR2-dependent manner since the blockade of TLR2 abrogates the CEP effect (Supplementary fig. 5D, 5E). In another model, CEP promotes cell proliferation of human hair follicle dermal papilla cells (HFDPC) by ∼2-fold compared to control (Supplementary fig. 5F).

Next, we show that CEP promotes hair regeneration in injury in a TLR2-dependent manner. CEP administration promoted HF regeneration in WT wounds. However, it was ineffective in global TLR2^KO^ mice (Supplementary fig. 5H). CEP promoted a 55% increase in the number of HF and cell proliferation in WT wounds, and at the same time, there was no effect in TLR2^KO^ wounds (Supplementary fig. 5I-5J).

To ensure independence from immune cells, WT and TLR2^KO^ mice were irradiated and transplanted with WT bone marrow prior to wounding. Applying CEP on wounds in WT/WT chimeras promoted cell proliferation, thereby dramatically increasing the density of hair follicles (Fig.6H, 6I, Supplementary fig. 5G). At the same time, CEP was not effective in TLR2^KO^/WT mice (Fig.6H, 6I, Supplementary fig. 5G), demonstrating the TLR2-dependent mechanism.

These CEP effects were mediated by TLR2 on HFSCs. In control mice, CEP effectively initiated regeneration of HFs around the wound (Fig.6J), resulting in ∼3-fold higher density of HF (Fig.6K, 6L) and dramatic acceleration of cell proliferation by >10 fold (Fig.6M, 6N) as compared to TLR2^HFSC-KO^ mice where CEP was mainly ineffective. A similar stimulatory effect of CEP was observed in a primary keratinocyte culture (Oshimori and Fuchs, 2012). CEP dramatically promoted WT but not TLR2^KO^ keratinocyte proliferation (Fig. 6O, 6P). CEP was an effective stimulator of TLR2 signaling as judged by augmented *Nfkb2*, *Il1β,* and *Il6* expression in HFSCs upon treatment with CEP (Fig. 6Q). Consistent with the key role of BMP signaling in TLR2-dependent HF regeneration, CEP treatment suppressed inhibitory *Bmp7* expression by ∼2.5 fold (Fig. 6R), demonstrating that endogenous and natural TLR2 ligand can counteract an inhibitory effect of BMP7 to stimulate HFSCs activation (Fig. 6S).

## Discussion

The main findings of this study are as follows: 1) Expression of TLR2 in HFSCs is decreased with aging and in a mouse model of obesity. 2) In young and healthy animals, TLR2 expression in HFs is cycle-dependent, with the highest expression in HFSCs during the initiation of the anagen phase. 3) The absence of TLR2 in HFSCs prolongs the resting phase of the hair cycle and significantly delays hair regeneration after injury. 4) TLR2 regulates the hair cycle primarily by inhibiting BMP signaling in HFSCs. 5) HFs continuously produce a metabolite of PUFAs, which acts as an endogenous TLR2 ligand and promotes hair growth through TLR2 activation in HFSCs. Besides reduced TLR2, aging is linked to low levels of its ligand in HFs. The stimulatory role of TLR2 signaling in HFs was demonstrated through both animal models and established human cell lines.

The lack of TLR2 appears to shift the balance between activating and inhibitory cues, leading to a resting phase that is approximately three times longer. This is a substantial impact considering that there are only 4-5 hair cycles in a mouse lifetime (Choi et al., 2021). The decrease in both TLR2 and its ligand observed in aging and associated conditions will inevitably impede the cyclic regeneration of hair follicles.

The immune system was shown to play a role in the activation of HFSCs, even in the absence of inflammation (Ali et al., 2017, Castellana et al., 2014, Pinho and Frenette, 2019). TLR2 expression increases at the onset of anagen when the immune response is reduced (Paus et al., 2003) and the hair follicle is most susceptible to pathogens. The upregulation of TLR2 in anagen may initially have a protective function. However, high TLR2 expression in undifferentiated vs. differentiated cells underscores its role in stem cell biology. TLRs, including TLR2, have been shown to play a critical role in stem cell functions in various organs (Lathia et al., 2008, Tomchuck et al., 2008, Trowbridge and Starczynowski, 2021). TLRs’ ligation and signaling can alter stem cell differentiation patterns (Collins et al., 2021, Nagai et al., 2006, Trowbridge and Starczynowski, 2021). Proinflammatory signaling can also activate HFSC proliferation, e.g., during injury (Wang et al., 2017, Chen et al., 2015). During the immune-privileged anagen phase of the hair cycle, TLR2 signaling may act as a key intrinsic factor in triggering HFSC activation.

The role of innate immunity in stem cell activation has mainly been linked to TLR3, shown to induce pluripotency in somatic cells through nuclear reprogramming (Lee et al., 2012, Sayed et al., 2015) and drive hair follicle (HF) neogenesis after tissue damage (Nelson et al., 2015). In contrast, we show that TLR2 drives a rapid inflammatory response and regulates the normal hair cycle and HF regeneration/neogenesis in injury.

TLR2 promotes the hair cycle by inhibiting the BMP pathway, a key regulator of hair follicle stem cell (HFSC) quiescence (Plikus et al., 2008, Hsu et al., 2011, Kandyba et al., 2013). Our study demonstrates that reducing excessive BMP signaling reactivates TLR2-deficient HFSCs, revealing a novel link between TLR2, BMP signaling, and the hair cycle. The only known instance of immune system-mediated BMP pathway inhibition occurs during the apoptosis of bulge-associated macrophages (Castellana et al., 2014, Wang et al., 2017).

The role of TLR2 in hair follicle regeneration in both normal hair growth and wound healing emphasizes the importance of understanding the nature of TLR2 ligands mediating these responses. At the site of injury, TLR2 can be activated by pathogens or by endogenously produced ligands, such as the oxidative product of PUFA, CEP, generated in abundance during wound healing (West et al., 2010, Yakubenko et al., 2018). CEP and TLR2 are both essential for hair regeneration, and their deficiency observed in pathologies such as aging and obesity might substantially impair hair growth. Exogenous application of CEP accelerates both wound closure (West et al., 2010) and HF regeneration through TLR2. In addition, both TLR2 and the application of CEP diminish inhibitory BMP signaling, suggesting that CEP and other TLR2 ligands could have therapeutic value for the treatment of hair loss related to burns, traumas, and other pathologies. It is intriguing that while CEP is almost exclusively generated at sites of injury and inflammation (Yakubenko et al., 2018, Xiong et al., 2022a), HFs continuously produce it, most likely by the means of MPO, an anti-bacterial enzyme, capable of generating CEP (Klebanoff et al., 2013).

Contrary to the trend observed in other tissues, where the accumulation of oxidation-generated CEP increases with aging (West et al., 2010), hair follicles (HFs) seem to exhibit depletion of CEP with aging. This decline in CEP levels might contribute to the reduced activity of hair follicle stem cells (Fuchs and Blau, 2020).

The role of CEP in TLR2-dependent HF growth and regeneration highlights the connection between oxidative stress and regenerative processes. Sustained reactive oxygen species (ROS) play a crucial role in proper regeneration, as seen in the tail amputation of Xenopus tadpole (Love et al., 2013). ROS enhance the differentiation of hematopoietic progenitors in Drosophila (Owusu-Ansah and Banerjee, 2009) and sustain self-renewal in neural stem cells (Le Belle et al., 2011). Additionally, ROS production in the skin has been linked to the activation of hair follicle stem cells (HFSCs) (Carrasco et al., 2015). We show the underlying mechanism for these observations, where oxidation-generated CEP triggers TLR2 activation, decreases inhibitory BMP signaling, and stimulates hair follicle growth and regeneration. TLR2 appears to serve as a common link between oxidative stress and tissue regeneration.

To summarize, our study highlights a novel role of TLR2 in promoting tissue regeneration during normal hair growth and wound healing. The identification of an endogenous TLR2 ligand produced by hair follicles presents a potential target for augmenting hair regeneration in the context of injury and aging, opening up new avenues for regenerative medicine.

## List of abbreviations

TLR: toll-like receptor
HFs: hair follicles
HFSCs: hair follicle stem cells
CEP: carboxyethylpyrrole
MPO: myeloperoxidase
PUFA: polyunsaturated fatty acid
sHG: secondary hair germ
DP: dermal papilla
ORS: outer root sheath
IRS: inner root sheath
MPPs: multipotent progenitors
NHEK: human epidermal keratinocytes
HFDPC: hair follicle dermal papilla cells

## Materials and Methods

### Mice

Inducible K15-CrePR1 mice (Stock No. 005249), TLR2-GFP reporter mice Stock No. 031822), and TLR2^KO^ mice (Stock No. 004650) were purchased from the Jackson Laboratory. TLR2^flox/flox^ mice with Exon3 of the *Tlr2* gene flanked by two loxP sites were described elsewhere (McCoy et al., 2021). HFSC-specific *Tlr2*-knockout (*Tlr2*^HFSC-KO^) mouse line was described previously (Xiong et al., 2022b). Briefly, TLR2^flox/flox^ mice were crossed with K15-CrePR1 mice to generate the inducible hair follicle stem cell-specific TLR2 knockout mouse line. To induce Cre recombinase activity, RU486 (Sigma) was used topically on shaved dorsal skin (1% mixed with Neutrogena Hand Cream) or via intraperitoneal injection (10 mg/mL in corn oil, 75ug RU486 per 1kg body weight) during the first postnatal telogen. To block BMP signaling in mouse hair follicles, at first postnatal telogen after applying Ru486, 200 ng of recombinant mouse Noggin (Biolegend) reconstituted in 30 µl of PBS were injected intradermally into a dorsal skin for 3 to 5 consecutive days. BSA in PBS was used as vehicle control.

For high-fat diet feeding studies, male wild type C57BL/6J were purchased from Jackson Laboratories (Bar Harbor, ME), and at seven weeks of age, mice were either maintained on standard rodent chow or switched to a high-fat diet containing 60% of kilocalories from fat (Research Diets D12492) for an additional 15 weeks prior to tissue collection. All procedures were performed according to animal protocols approved by the Cleveland Clinic IACUC committee.

### Immunostaining

Mouse skin samples were harvested at indicated ages and fixed in 4% paraformaldehyde, kept in 30% sucrose for two to three days, followed by snap-freezing at -80℃ in OCT (Fisher HealthCare, 4585). 10um skin sections were permeabilized, blocked, and incubated with primary antibodies followed by incubation with the corresponding secondary antibody, and mounted with an antifade mounting medium with DAPI (Vector Laboratories, H-1500-10). Images were captured on a Leica DM2500 confocal microscope and analyzed using Bitplane Imaris software (version 9.7.2) or ImageJ. Briefly, Image Z-stacks were loaded into Imaris to reconstruct three-dimensional images, and surface rendering was performed with default settings using the surface tool. The same background subtraction was performed on each z-stack. The green channel was used as a source channel to create surfaces for GFP^+^ cells in the area of interest in hair follicles, and other channels were created based on the expression of different cell markers (e.g., CD34, Ker5) in the hair follicles. The overlap between GFP surface and other maker surfaces was created and visualized as the colocalized area with the Colocalization module. At least 10 hair follicles from each mouse were used for quantification.

The following antibodies or reagents were used: Ker17 (Santa Cruz Biotechnology, sc-393002), Ker15 (ABclonal, A2660), myeloperoxidase (Santa Cruz Biotechnology, sc-390109), GFP (Thermofisher Scientific, CAB4211), TLR2 (Santa Cruz Biotechnology, sc-21759), Ki67 (Abcam, ab16667), P-cadherin (R&D systems, AF761-SP), CEP (Pacific Immunology), pSmad1/5/9 (Cell Signaling Technology, 13820S,), β-catenin (Cell signaling Technology, 9581), CD34 (eBioscience, 11-0341-82), CD49f (BD Pharmigen, 562473), Ker5 (Biolegend, 905903), Sox9 (Cell Signaling Technology, 82630T), BMP7 (Proteintech, 12221-1-AP), and Nile Red (ATT BioQuest, 250730). As a negative control, we used appropriate isotype match nonimmune antibody: normal mouse IgG2b-PE (Santa Cruz Biotechnology, sc-2868), Normal Goat IgG Control (R&D, AB-108-C), normal mouse IgG (Santa Cruz Biotechnology, sc-2025), normal Rabbit IgG (Cell signaling Technology, 2729S).

### Isolation of keratinocytes and hair follicle stem cells

Keratinocytes and hair follicle stem cells were isolated from the mouse dorsal skin as described previously (Xiong et al., 2022b). Briefly, isolated dorsal skin samples were trypsinized, the epidermis was scrapped, minced, and filtered through a 70-µm cell strainer to prepare primary keratinocytes single-cell suspension. To isolate HFSC, the single-cell suspension was incubated with CD34-FITC antibody (eBioscience, 11-0341-82), Alexa 647-conjugated CD49f antibody (BD Biosciences, 562494), 7-AAD (BD Biosciences, 559925), and different Fluorescence minus one was used as a control. Cells then were sorted by BD FACS Aria and analyzed by Flow Jo.

### Wound healing

Mouse wound healing procedure was performed as previously described (West et al., 2010, Xiong et al., 2022b). Briefly, an intraperitoneal injection of a ketamine/xylazine cocktail was used to anesthetize 7-to 8-week-old mice. After shaving, full-thickness wounds were made into the dorsal skin using a 6mm biopsy punch. To examine the effect of CEP on hair regeneration after wound healing, CEP (CEP in polyethylene glycol) or vehicle (polyethylene glycol) was applied to the wounded area every day for two weeks. Pictures were taken at different time points to record hair regeneration around the wounded area.

### Primary keratinocyte proliferation assay

The primary keratinocytes after isolation were plated on rat tail collagen-coated plates with Epilife medium (Gibco, MEPI500CA) supplemented with EDGS (Gibco, S0125). Cells were co-cultured with CEP or control (BSA or PBS) for 48 hours. The cell counting kit 8 (APEXbio, 269070) has been used to measure cell proliferation according to the manufacturer’s protocol.

### Human Hair Follicle Dermal Papilla Cells proliferation assay

Human Hair Follicle Dermal Papilla Cells (HFDPC) (Cell applications inc. cat.# 602-05a) were cultured in HFDPC Growth Medium (Cell applications inc. cat.#611-500) for 60 hours and then transferred into collagen-coated 48 wells plate for 24 hours. After 24 hours cells were washed with 1x D-PBS and incubated in HFDPC Basal Medium contains no growth supplement (Cell applications inc. cat.#610-500) for the next 24 hours. After starvation, cells were incubated with CEP 5 µM or with HFDPC Growth Medium (positive control), or in HFDPC Basal Medium (negative control) for 48 hours. Absorbance was read using Cell counting kit 8 (ApexBio, cat.# K1018) on a microplate reader.

### Human Hair Follicle Stem Cells proliferation assay

Human Hair Follicle Stem Cells (HFSC) (Celprogen cat.# 36007-08) were cultured in HFSC Un-differentiation Media with Serum (Celprogen cat.# M36007-08US) for 48 hours and then transferred into Undifferentiated ECM -96 Well Plates (Celprogen cat.# UD36007-08-96Well) for 24 hours. After 24 hours cells were washed with 1x D-PBS and incubated in HFSC Serum Free Un-differentiation Media (Celprogen cat.# M36007-08U) overnight. After starvation, cells were incubated for 2 hours with or without TLR2 blocking antibody (Invivogen cat.# mab2-mtlr2) followed by incubation with CEP or Pam3CSK4 (Invivogen cat.# tlrl-pms) for 24 hours. HFSC Serum Free Un-differentiation Media was used as a negative control. Absorbance was read using Cell counting kit 8 (ApexBio, cat.# K1018) on a microplate reader.

### Human Epidermal Keratinocytes experiments

Human Epidermal Keratinocytes (Lonza Reagents cat#192906) were cultured in KGMTM Gold Keratinocyte Growth Medium (Lonza Reagents cat#192060) for 60 hours and then transferred into a 6-wells plate for 24 hours. After 24 hours media was changed and cells were incubated with Pam3CSK4 10 µg /ml for 1 hour followed by incubation with BMP4 10 ng/ml for 1 hour.

### CEP synthesis and preparation

The structure and synthesis of CEP (carboxyethylpyrrole) have been described elsewhere (West et al., 2010). To prepare CEP for wound healing, 250uL CEP in PBS was mixed with 1.1g polyethylene glycol with sonication in a 45℃ water bath for 15 minutes followed by a strong vortex to mix well. This mixture was stored at 4℃ after preparation, warmed to room temperature, and mixed again before use.

### Real-time qPCR

Total RNA from primary keratinocytes or hair follicle stem cells was isolated with RNeasy Mini Kit (Qiagen, 74104) and reverse transcribed into cDNA with PrimeScript RT Master Mix (Takara, RR036A). The real-time polymerase chain reaction was performed using iQ SYBR Green Supermix (Bio-Rad, 1708882) on the Bio-Rad cfx96 qPCR system. Target gene expression levels were normalized to internal control Rps16, and the ΔΔCt method was used to calculate fold change in gene expression. Primers can be found in Table 1.

### Western blot analysis

Cells were lysed with RIPA Lysis and Extraction Buffer (ThermoScientific™cat.#PI89900) buffer with protease/phosphatase inhibitor cocktail. The lysate was centrifuged at 12000g at 4^0^C for 15 min, boiled with Laemmli buffer for 7 min at 95^0^C, and transferred to PVDF membranes (Millipore). After blocking, membranes were incubated with primary antibody at 4^0^C overnight followed by incubation with corresponding secondary HRP-linked antibody. The following antibodies were used for western blotting: Smad1 (D59D7) XP® Rabbit mAb (Cell Signaling Technology cat#6944), Phospho-Smad1 (Ser463/465)/ Smad5 (Ser463/465)/ Smad9 (Ser465/467) (Cell Signaling Technology cat# CLS8160-24EA), NF-κB p65 (D14E12) XP® Rabbit mAb (Cell Signaling Technology cat#8242), Phospho-NF-κB p65 (Ser536) (93H1) Rabbit mAb (Cell Signaling Technology cat#3033), Anti-GAPDH antibody EPR16884 Loading Control (Abcam cat#ab181603).

### Bone marrow transplant (BMT) and wound assay

We performed BMT as previously described (West et al., 2010) Briefly, two-month-old male WT or TLR2^KO^ mice were lethally irradiated with 9 Gy followed by tail vein injection with 10^7^ bone marrow cells isolated from the WT donor femurs. Eight weeks after BMT, mice were subjected to wound healing assay (described above).

### RNA sequencing and data analysis

First telogen mouse dorsal skin was used for hair follicle stem cell isolation by FACS. Total RNA was extracted using the RNeasy Mini Kit (Qiagen, 74104). Sample quality assessment was performed on a Fragment Analyzer electrophoresis system (Agilent). Total RNA was normalized prior to oligo-dT capture and cDNA synthesis with SMART-Seq v4 (Takara). The resulting cDNA was quantified using a Qubit 3.0 fluorometer (Life Technologies). Libraries were generated using the Nextera XT DNA Library Prep kit (Illumina). Medium-depth sequencing (50 million reads per sample) was performed with a NextSeq 550 (Illumina) on a High Output flow cell using 75 base pairs, Paired-End run. Raw demultiplexed fastq paired-end read files were trimmed of adapters and filtered using the program skewer to throw out any with an average Phred quality score of less than 30 or a length of less than 36. Trimmed reads were then aligned using the HISAT2 aligner to the Mouse NCBI reference genome assembly version GRCm38 and sorted using SAMtools. Aligned reads were counted and assigned to gene meta-features using the program featureCounts as part of the Subread package. These count files were imported into the R programming language and were assessed for quality control, normalized, and analyzed using an in-house pipeline utilizing the limma-trend method for differential gene expression testing and the GSVA library for gene set variation analysis. The pathway analysis for differentially expressed genes with adjP-value<0.05 was performed using Enrichr web server https://maayanlab.cloud/Enrichr

#### Statistical analysis

Statistical analyses were performed using GraphPad Prism 9. All results are mean ± s.e.m. Shapiro-Wilk normality and lognormality test was used with n≥6. For normally distributed data, we use an unpaired two-tailed t-test to compare two groups and the one-way ANOVA followed by Dunnett’s or Tukey’s post hoc analysis to compare more than two groups. For non-normally distributed data and small sample size (n<6), we appraised statistical differences with the non-parametric Mann-Whitney test to compare two sample data sets and the Kruskal-Wallis test with Dunn’s post hoc test for 3 and more groups. A p-value < 0.05 was considered to be statistically significant.

## Acknowledgments

We thank D. Nascimento, and K. Li for mouse colony management; T. Dudiki for revision of figures; J. Powers for FACS assistance; C. Nelson for proofreading. Applied Functional Genomics Core for RNA sequencing and M. Kumar for data analysis.

## Author Contributions

Conceptualization: TVB

Methodology: LX, XW, IZ, RM, DG, TVB

Investigation: LX, XW, IZ, RM, IM, TVB

Data curation: LX, IZ, XW, RM, TVB

Funding acquisition: TVB

Resources: DG, EP, AH, TVB

Supervision: TVB, EP.

Writing – original draft: LX, IZ, TVB

Writing – review & editing: JMB, TVB

## Data availability

The RNAseq dataset is available in the Gene Expression Omnibus GSE179300

