## Supplemental data for "TLR2 Regulates Hair Follicle Cycle and Regeneration via BMP Signaling"

**Supplemental material.**

**Supplemental figure 1.** TLR2/GFP correlation in immunostaining. H&E staining of dorsal skin of WT mice. TLR2 dynamic during the second telogen.

**Supplemental figure 2.** Confocal images of hair follicles immunostained for P-cad

**Supplemental figure 3.** Bulge stem cell marker expression in $TLR2^{HFSC\text{-}KO}$ mouse.

**Supplemental figure 4.** TLR2-BMP axis in hair follicle cells. BMP signaling in the hair bulge of WT and $TLR2^{HFSC\text{-}KO}$ mice.

**Supplemental figure 5.** MPO expression in sebaceous gland of hair follicles from old vs young mice; effect of CEP in-vitro and in-vivo on hair follicle growth.

**Supplemental table 1.** qPCR primers.

**Supplementary figure 1**

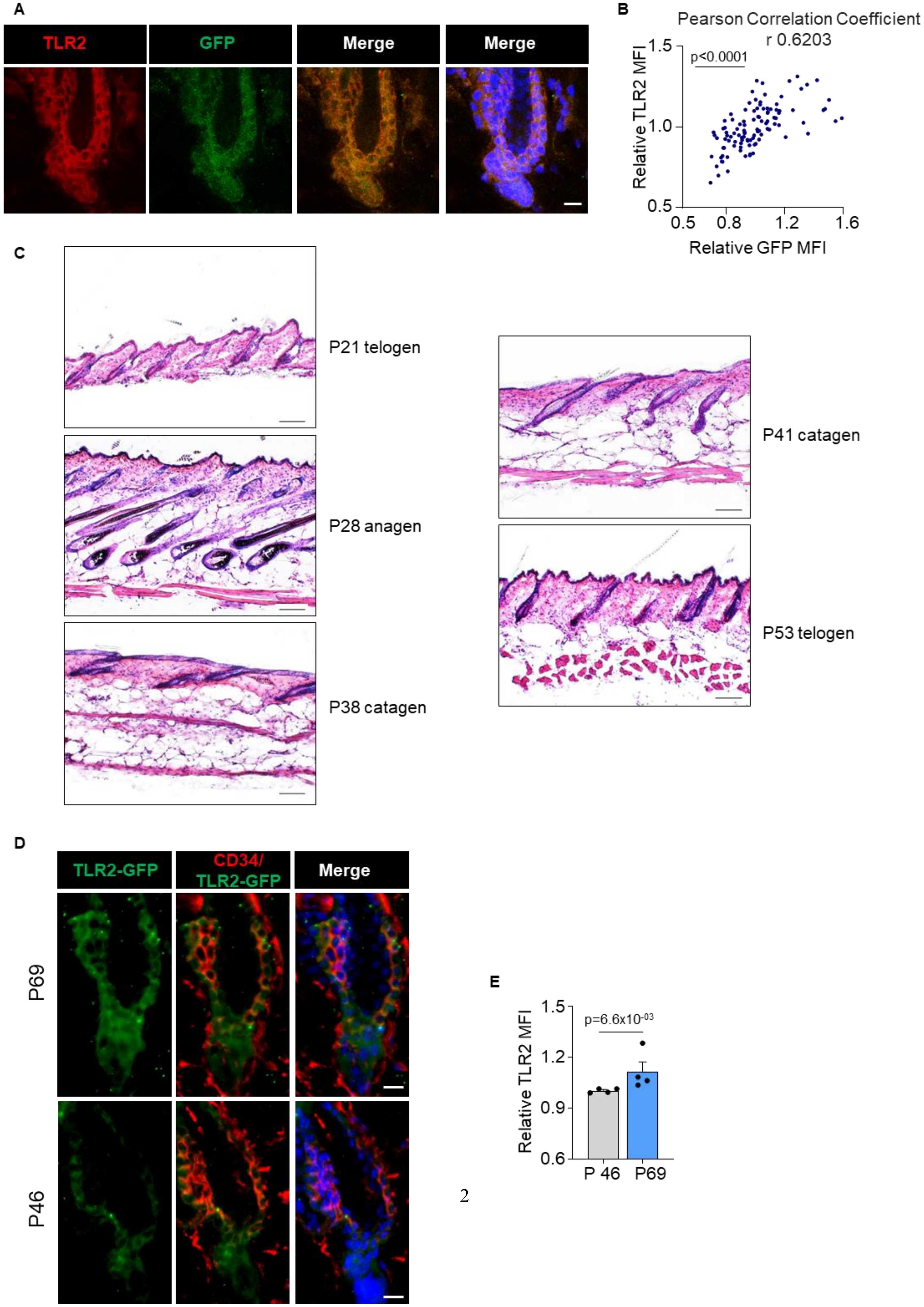

**Fig. S1.**

**A.** Confocal images of dorsal skin hair follicles co-immunostained for TLR2 and GFP. Scale bars are 10 µm.

**B**. A scatter graph shows a high level of correlation between TLR2 and GFP intensity. N=3.

**C.** Representative H&E staining of the dorsal skin of WT mice at indicated time points demonstrates the typical changes in hair follicle morphology. Scale bars are 100 µm.

**D**. Confocal images of dorsal skin hair follicles from p46 and p69 mice immunostained for TLR2.

**E.** Bar graph showing an elevated level of TLR2 in HFSC of p69 mice (late second telogen) compared to p46 (early second telogen). N = 4

Correlation analysis and the Mann-Whitney test were used to determine the statistical significance. All data are mean ± s.e.m.

**Supplementary figure 2**

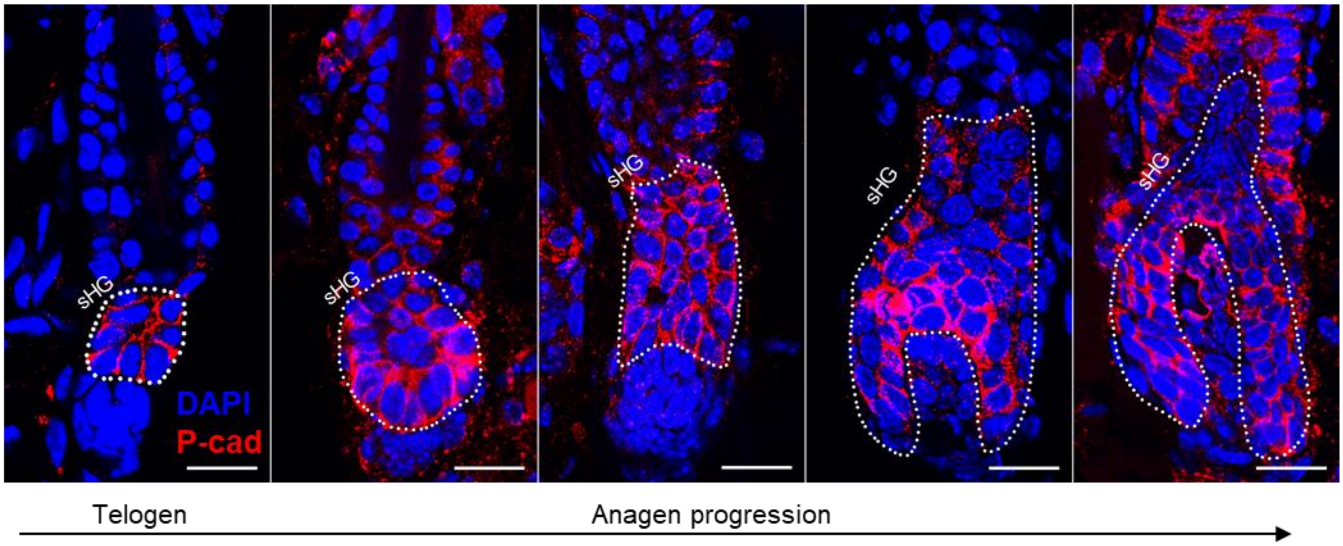


**Fig.S2.**

**sHG enlargement and elongation at anagen onset.**

Representative confocal images of hair follicles immunostained for P-cad showing changes in the size of sHG during the progression of the hair cycle. At anagen onset, sHG cells proliferate and grow downward to envelop the dermal papilla. The proliferation of sHG cells results in a larger sHG compared to telogen. Dashed lines outline the sHG. Scale bars are 20 μm.

**Supplementary figure 3**

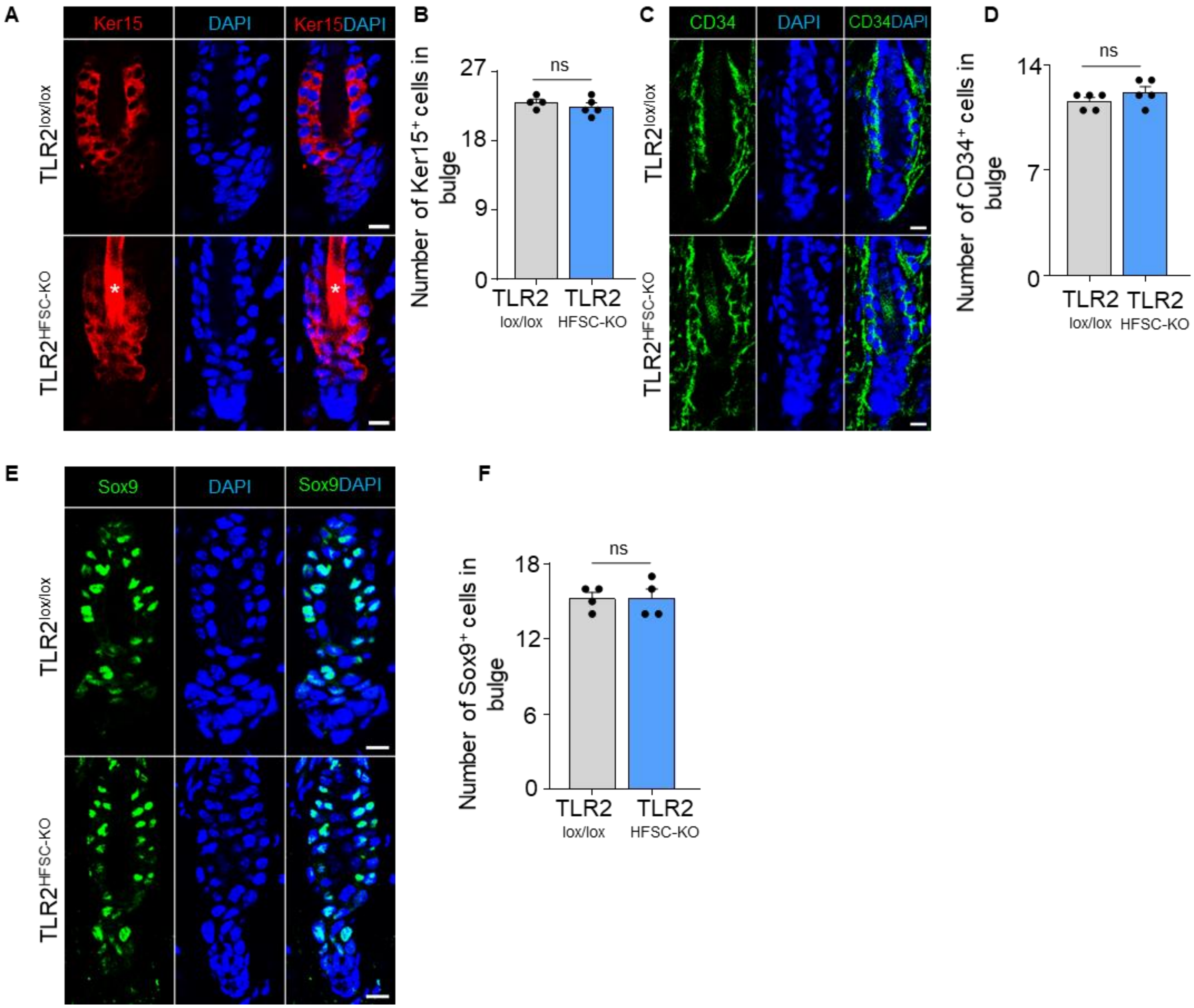

**Fig.S3.**

**Bulge stem cell marker expression in TLR2$^{HFSC-KO}$ mouse.**

**A.** Representative confocal images of telogen hair follicles from TLR2$^{lox/lox}$ and TLR2$^{HFSC-KO}$ mice immunostained for Ker15. Stars label the hair shaft. Scale bars are 10 µm.

**B.** Bar graph showing no difference in numbers of Ker15$^{+}$ cells in bulge area in TLR2$^{lox/lox}$ compared with TLR2$^{HFSC-KO}$ telogen hair follicles from images in A. N = 4 and 5 mice for TLR2$^{lox/lox}$ and TLR2$^{HFSC-KO}$ respectively.

**C.** Telogen hair follicles from TLR2$^{lox/lox}$ and TLR2$^{HFSC-KO}$ mice immunostained for CD34. Scale bars are 10 µm.

**D.** Quantification of CD34$^{+}$ cell numbers in bulge area in images from C showing no difference in TLR2$^{lox/lox}$ compared with TLR2$^{HFSC-KO}$ follicles. N = 5 per group.

**E.** Representative confocal images of TLR2$^{lox/lox}$ and TLR2$^{HFSC-KO}$ telogen hair follicles immunostained for Sox9. Scale bars are 10 µm.

**F.** Bar graph showing no difference in Sox9$^{+}$ cell numbers in bulge area in TLR2$^{lox/lox}$ and TLR2$^{HFSC-KO}$ follicles from images in E. N = 4.

Mann-Whitney test was used to determine the statistical significance. All data are mean ± s.e.m.

#### Supplementary figure 4

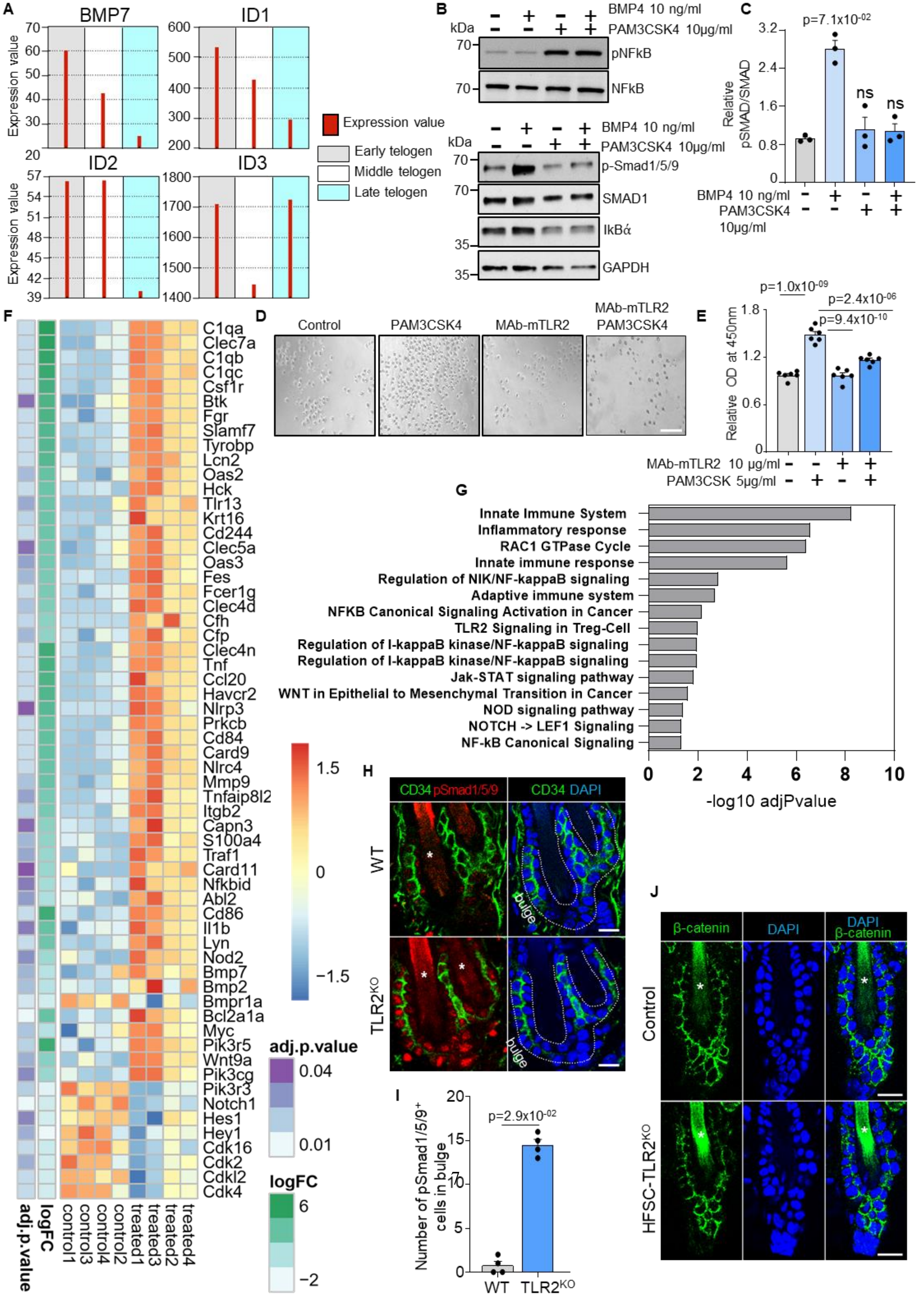

**Fig. S4.**

**A.** GEO2R analysis of published RNA data from sorted follicle populations in the 2nd telogen to anagen transition demonstrates the declined expression of BMP7 mRNA and key transcriptional target of the BMP signaling pathway in the skin, ID1, ID2, and ID3 mRNA.

**B.** Western blot analysis of pNFkB, NFkB, pSMAD1/5/9, SMAD1, ikBα protein level in Human Epidermal Keratinocytes treated with/without 10 μg/ml Pam3SCK4 and 10 ng/ml BMP4.

**C.** Bar graphs show the activation effect of BMP4 treatment on the BMP signaling which was abolished in the presents of TLR2 ligand Pam3SCK4. N=3 independent experiments.

**D.** Representative microphotographs of Human Hair Follicle Stem Cells pre-treated with 10 μg/ml MAb-mTLR2 or DMSO and co-cultured with/without 5μg/ml Pam3SCK4. Representative images from at least three independent assays are shown. Scale bar 50 μm.

**E.** Bar graphs show increased proliferation of HFSC in the presents of TLR2 ligand Pam3SCK4 with no effect in MAb-mTLR2 pre-treated cells compared to control. N=6 independent experiments.

**F.** Dysregulated genes were confirmed by RNA sequencing or qPCR of FACS-sorted HFSCs from the first telogen dorsal skin of $TLR2lox^{/lox}$ and $TLR2^{HFSC-KO}$ mice. HFSCs from 4 mice in each group were sorted and sequenced. Dysregulated genes in RNA sequencing were those with an adjusted p-value < 0.05.

**G.** Dysregulated pathways in HFSCs lacking TLR2.

**H.** Representative confocal images of competent telogen follicles from WT or $TLR2^{KO}$ mice immunostained for CD34 and pSmad1/5/9. Stars label hair shaft. Scale bars are 10 μm.

**I.** Quantification of $pSmad1/5/9^{+}$ bulge stem cell number in images from H shows significantly more $pSmad1/5/9^{+}$ cells in WT hair follicle bulge region compared with $TLR2^{KO}$ hair follicles. N = 4 for each group**.**

**J**. Representative confocal images of $TLR2^{lox/lox}$ and $TLR2^{HFSC-KO}$ telogen hair follicles immunostained for β-catenin showed no difference between the two groups. Stars label the hair shaft. Scale bars are 20 μm.

All bar graphs are mean ± s.e.m. Non-parametric Mann-Whitney test (I), or Kruskal-Wallis test with Dunn's multiple comparisons test (C), or one-way ANOVA with Tukey's multiple comparisons test (E) was used to determine statistical differences. For the WB quantification with experiments with n=3, the minimum achievable p-value for the non-parametric tests is 0.1000.

### Supplementary figure 5

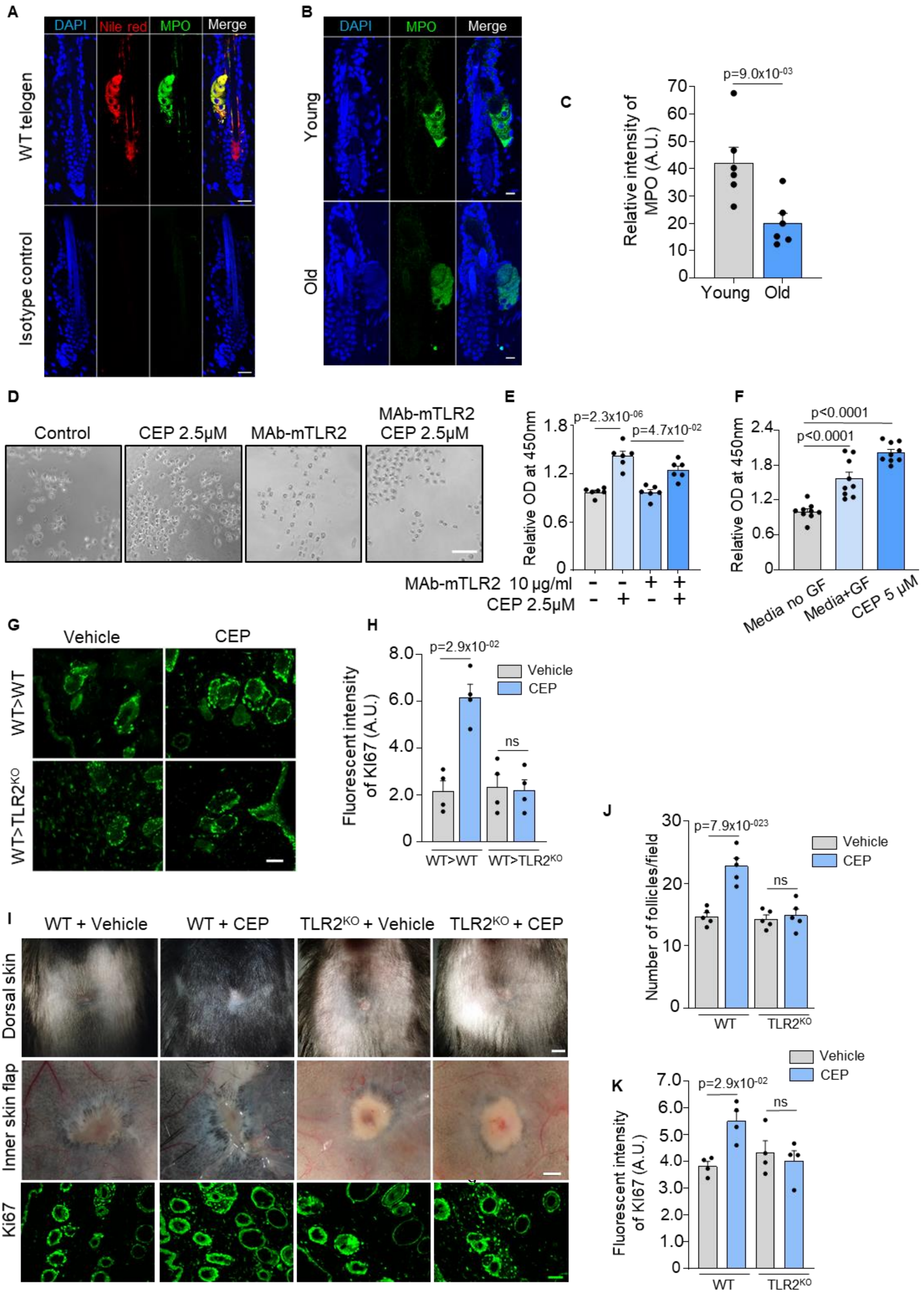

**Fig. S5.**

**The promotion of hair follicle regeneration after wound healing is dependent on TLR2**

**A.** Representative confocal images of Nile red-labeled (sebaceous gland) WT telogen hair follicles co-immunostained for myeloperoxidase (MPO) showing complete co-localization of MPO to the sebaceous gland. The isotype control panel shows the images of hair follicles stained with MPO isotype control antibody. Scale bars are 20 µm.

**B.** Representative confocal images of hair follicles from old vs young mice stained for MPO. Scale bars are 10 µm.

**C.** Quantification of MPO fluorescent intensity in B showing significantly less MPO in hair follicles from older mice. N = 6 for each group.

**D**. Representative microphotographs of Human Hair Follicle Stem Cells pre-treated with 10 µg/ml MAb-mTLR2 or DMSO and co-cultured with/without 2.5µM of CEP. Representative images from at least three independent assays are shown. Scale bar 50 µm.

**E**. Bar graphs show increased proliferation of HFSC in the presence of TLR2 endogenous ligand CEP compared to control, which was abolished in the presence of TLR2 blocking antibody. N=6 independent experiments.

**F.** Bar graphs show increased proliferation of Human Hair Follicle Dermal Papilla Cells incubated with 5µM of CEP compared to the control. N=9 independent experiments.

**G**. Representative confocal images of Ki67 immunostaining of dorsal skin adjacent to wound of CEP-treated WT bone-marrow transplanted WT and $TLR2^{KO}$ mice. Scale bars are 50 µm.

**H**. Quantitative results showed increased Ki67 intensity in hair follicles around wounds of CEP-treated WT bone-marrow transplanted WT mice with no differences in $TLR2^{KO}$ with WT bone marrow. N = 4 per group.

**I.** Representative photographs of dorsal skin (upper panels), inner skin flaps (middle panels), and representative confocal images of Ki67 immunostaining (lower panels) of vehicle- or CEP-treated WT or $TLR2^{KO}$ skin. Scale bars are 1 mm for dorsal skin, 500 µm for skin flaps, and 50 µm for confocal images.

**J.** Bar graph showing quantification of hair follicle numbers of vehicle or CEP-treated skin from I. N = 5 per group.

**K.** Bar graph showing quantification of Ki67 fluorescent intensity of Ki67 staining of vehicle or CEP-treated skin from I. N = 4 per group.

Unpaired two-tailed t-test (C), or non-parametric Mann-Whitney test (H, J, K), or Kruskal-Wallis test with Dunn's multiple comparisons test (F), or one-way ANOVA with Tukey's multiple comparisons test (E) was used to determine statistical differences. All bar graphs are mean ± s.e.m.

**Table S1.**

| Target | Primers |
|---|---|
| *Bmp7* | ACGGACAGGGCTTCTCCTAC |
| | ATGGTGGTATCGAGGGTGGAA |
| *Bmp2* | GGGACCCGCTGTCTTCTAGT |
| | TCAACTCAAATTCGCTGAGGAC |
| *Bmpr1a* | AACAGCGATGAATGTCTTCGAG |
| | GTCTGGAGGCTGGATTATGGG |
| *Nfkb2* | GGCCGGAAGACCTATCCTACT |
| | CTACAGACACAGCGCACACT |
| *IL1b* | GCAACTGTTCCTGAACTCAACT |
| | ATCTTTTGGGGTCCGTCAACT |
| *IL6* | TAGTCCTTCCTACCCCAATTTCC |
| | TTGGTCCTTAGCCACTCCTTC |
| *Hey1* | GCGCGGACGAGAATGGAAA |
| | TCAGGTGATCCACAGTCATCTG |
| *Notch1* | ACACTGACCAACAAATGGAGG |
| | GTGCTGAGGCAAGGATTGGA |
| *Hes1* | CCAGCCAGTGTCAACACGA |
| | AATGCCGGGAGCTATCTTTCT |
| *Bcl2a1a* | GGCTGAGCACTACCTTCAGTA |
| | TGGCGGTATCTATGGATTCCAC |
| *Myc* | ATGCCCCTCAACGTGAACTTC |
| | CGCAACATAGGATGGAGAGCA |
| *Tlr2* | TCTAAAGTCGATCCGCGACAT |
| | CTACGGGCAGTGGTGAAAACT |

qPCR primers.
